# Decelerated carbon cycling by ectomycorrhizal fungi is controlled by substrate quality and community composition

**DOI:** 10.1101/716555

**Authors:** Christopher W. Fernandez, Craig R. See, Peter G. Kennedy

**Affiliations:** Department of Plant and Microbial Biology, University of Minnesota; Department of Ecology, Evolution and Behavior, University of Minnesota

**Author notes:** Corresponding author, Contact Information: 1479 Gortner Ave., St. Paul, MN, 55108.

**Keywords:** carbon, competition, decomposition, ectomycorrhizal fungi, Gadgil effect, litter, nitrogen cycle, saprotrophic fungi, soil organic matter

## Abstract

Interactions between symbiotic ectomycorrhizal (EM) and free-living saprotrophs can result in significant deceleration of leaf litter decomposition. While this phenomenon is widely cited, its generality remains unclear, as both the direction and magnitude of EM fungal effects on leaf litter decomposition have been shown to vary among studies. Here we explicitly examine how contrasting leaf litter types and EM fungal communities may lead to differential effects on C and N cycling. Specifically, we measured the response of soil nutrient cycling, litter decay rates, litter chemistry and fungal community structure to the reduction of EM fungi (via trenching) with a reciprocal litter transplant experiment in adjacent *Pinus*- or *Quercus*-dominated sites. We found clear evidence of EM fungal suppression of C and N cycling in the *Pinus*-dominated site, but no suppression in the *Quercus*-dominated site. Additionally, in the *Pinus*-dominated site, only the *Pinus* litter decay rates were decelerated by EM fungi and were associated with decoupling of litter C and N cycling. Our results support the hypothesis that EM fungi can decelerate C cycling via N competition, but strongly suggest that the ‘Gadgil effect’ is dependent on both substrate quality and EM fungal community composition. We argue that understanding tree host traits as well as EM fungal functional diversity is critical to a more mechanistic understanding of how EM fungi mediate forest soil biogeochemical cycling.

## Introduction

Soil organic matter (SOM) decomposition is a critical nexus in the global cycling of carbon (C) and nitrogen (N), and a flux with cascading effects on a range of important ecosystem services, including nutrient availability and soil C stabilization (Schlesinger and Bernhardt, 2013). In forests, soil fungi are the primary agents of decomposition through the production of extracellular enzymes that break down SOM to acquire growth-limiting resources (Baldrian, 2017). The two dominant fungal guilds involved in forest soil SOM decomposition in are free-living saprotrophic fungi and symbiotic ectomycorrhizal (EM) fungi (Lindahl & Tunlid, 2015). These fungi potentially compete with each other as well as other soil biota for resources found in SOM. Unlike saprotrophs, however, EM fungi are not limited by the C in SOM, since it is provided by their tree hosts in the form of simple sugars (Smith & Read, 2008). This is thought to allow EM fungi to allocate resources towards exploiting soil nutrient patches, particularly N, which can be scarce in temperate and boreal forest soils (Kaye & Hart, 1997). The resultant ‘N mining’ (Kuyper, 2017) by EM fungi would increase the C:N ratio of SOM substrates, limiting saprotrophic growth as those decomposers become increasingly N limited. This scenario creates a positive feedback loop, ultimately resulting in the accumulation of C stored in soil SOM (Gadgil & Gadgil, 1971). This phenomenon, referred to as the ‘Gadgil effect’, has received renewed interest due to the potential of soil C storage to counteract increases in atmospheric CO2 concentrations (Orwin et al., 2011; Averill & Hawkes, 2016).

Despite widespread reference to this phenomenon in the literature, knowledge about the generality of the ‘Gadgil effect’ remains limited (Fernandez & Kennedy, 2016). In particular, field-based experiments implementing EM reduction treatments (e.g. soil trenching and tree girdling) in different forest systems have generated inconsistent results, calling into question the ubiquity of this phenomenon. One of the most important yet poorly understood biotic factors that may modulate the direction and magnitude of the ‘Gadgil effect’ is soil fungal community composition. With methodological advances in characterizing fungal communities (Nilsson et al. 2018), examining the effects of specific EM fungal and soil saprotrophic taxa may provide greater insight into how fungal-fungal interactions mediate rates of SOM decomposition. For example, there is growing evidence that members of the EM genus *Cortinarius*, which possess a range of class-II peroxidase genes, can significantly alter soil C stocks (Bödeker et al., 2014; Kyaschenko et al., 2017; Sterkenberg et al., 2018). Additionally, many EM fungal species are strongly host-specific, and different tree species in close proximity often have dramatically different associated EM fungal communities (Ishida et al., 2007; Tedersoo et al., 2008, Walker et al., 2014). Because of the functional diversity among EM fungi, particularly in their ability to explore and breakdown SOM (Kohler et al., 2015; Pellitier and Zak, 2017), variation in the composition of these communities may have strongly contrasting effects on forest C and N cycling (Zak et al., 2019). Similarly, taxonomic and functional variation among saprotrophic fungi may also influence the direction and magnitude of the ‘Gadgil effect’ (Van der Wal et al., 2013).

Along with differences in fungal community composition, variation in substrate chemistry is another key biotic variable that may influence how EM fungi affect SOM decomposition rates. Litter chemistry varies considerably across tree species (Berg 2000; Hobbie, 2008; Phillips et al., 2013), and both local and global analyses indicate that multiple chemical components are closely linked to decomposition rate, particularly N and lignin (Melilo & Aber 1982; Hobbie 2005; Cornwell et al., 2008). Since competition for N is the most commonly cited mechanism by which EM fungi may suppress saprotrophic decomposition (see Fernandez & Kennedy, 2016 for a discussion of alternative mechanisms), SOM with low N content may be particularly susceptible to this effect. Consistent with this prediction, EM fungal suppression of litter decomposition has been most pronounced in systems dominated by conifer trees (Fernandez & Kennedy, 2016), which typically have lower litter N content than angiosperm trees (Hobbie, 2005). In addition, there is growing evidence of ‘home field advantage’ (HFA) effects (Gholz et al., 2000), where litter decomposition is enhanced when litter and canopy composition are matched (Austin et al., 2014, Midgely et al., 2015). The mechanism(s) for HFA effects likely involve more than just optimization of microbial communities for specific substrates, as factors such as litter nutrient content alone do not capture their full magnitude (Vivanco & Austin, 2008). In fact, competitive interactions among different microbial guilds have been suggested to be important mediators of HFA effects (Van Der Wal et al., 2013). Although there has been some effort to decouple the effects of litter type and fungal interactions in tropical forests (McGuire et al. 2010), no studies to date have varied litter type and exclusion of EM fungi in the higher latitude forests where a ‘Gadgil effect’ has been most consistently observed.

The objective of this study was to understand how contrasting host tree and associated EM fungal communities may lead to differential effects on leaf litter decomposition and soil N cycling. To achieve this, we conducted two litter-bag decomposition experiments employing a soil trenching treatment, which disrupts host C flow into the plots and reduces EM fungal in-growth, in adjacent *Pinus* and *Quercus*-dominated forest sites in Minnesota, USA. In the first experiment, we designed a litter-bag decomposition study to establish the presence and consistency of the ‘Gadgil effect’ in our study system. In the following experiment, we used a reciprocal transplant design where both *Pinus* and *Quercus* litter were independently incubated in the same EM trenching treatments in *Pinus* and *Quercus*-dominated forests to assess if potential interactions between fungal community structure and litter type may ultimately govern decomposition dynamics. The composition of the entire fungal community colonizing the incubated litters in untrenched (control) and trenched plots was assessed after 2, 4, and 12 months with high-throughput sequencing. At each of those sampling times, changes in litter mass were measured and at 12 months, the C and N content of the incubated litter as well as inorganic soil N and P availability were assessed. Based on widespread presumption in the ‘Gadgil effect’ literature (e.g. Orwin et al., 2011; Averill et al. 2014), we hypothesized that EM suppression of litter decay rates would be associated with reductions in litter N content in plots where EM fungi were abundant. Based on hypotheses proposed in Fernandez & Kennedy (2016) and more recent modeling work by Smith and Wan (2019), we further hypothesized that given the lower initial N content of *Pinus* litter, that a ‘Gadgil effect’ would be more pronounced in this litter type than in *Quercus* litter. Finally, we speculated that inhibition of litter decomposition would occur primarily in association with EM fungal taxa known to produce enzymes associated with SOM decomposition.

## Materials & Methods

### Experimental design

Both experiments were conducted at the Cedar Creek Ecosystem Science Reserve in east-central Minnesota, USA. We located plots in two sites based on tree host composition, one being dominated by Northern pin oak (*Quercus ellipsoidalis*) (N 45.42142 W 093.19509) (hereafter referred to as the Oak site) and the other being dominated by Eastern white pine (*Pinus strobus*) (N 45.42577 W 093.20852) (hereafter referred to as the Pine site) (Table S1). These sites are *ca.* 1 km apart and have the same underlying sandy poorly developed Udipsamment soils, which have comparable soil pH and inorganic nitrogen concentrations within each soil layer (Table S2). On June 8, 2015, 6 randomly located blocks were established in each site, each containing a untrenched and a trenched EM fungal reduction treatment. The blocks were located at least 8 m apart to avoid spatial autocorrelation in fungal community composition (Lilleskov et al., 2004, Bahram et al., 2012). Trenching was done using a spade and cutting to a depth of 30 cm, which severed root and EM fungal in-growth into the plots. Given the newly generated inputs of labile carbon substrates (i.e. dead roots and fungal mycelium), the trenched plots were allowed to equilibrate after the initial disturbance for 5 weeks before commencing with the first experiment. To reduce root and EM fungal growth into the trenched plots, we carefully re-ran the spade through the trench slits on a bi-weekly basis during the growing season (July-November 2015, April-November 2016, April-July 2017). We measured the effect of the trenching on mineral soil moisture by taking soil cores (the same used later in molecular analyses) from each plot and determining the gravimetric water content on 5 g subsamples. We also monitored the effect of the trenching treatment on O-layer moisture content at five time points in the first growing season (July-November 2015) by collecting O-layer material and determining gravimetric moisture content from adjacent untrenched and trenched plots established for this purpose.

### Litter decomposition

Because forest soils at Cedar Creek are generally composed of unfragmented leaf litter directly above the soil A-layer, we used leaf litter as our substrate source, matching the original ‘Gadgil effect’ experiment (Gadgil & Gadgil, 1971) and the majority of studies that followed (Gadgil & Gadgil, 1975; Berg & Lindberger, 1980; Staaf, 1988; Zhu & Ehrenfeld, 1996; Koide & Wu, 2003; Mayor & Henkel, 2006; McGuire et al., 2010; Brzostek et al., 2015; Sterkenberg et al., 2018).

Recently senesced *Pinus strobus* and *Quercus ellipsoidalis* leaf litter (hereafter referred to as pine and oak litter, respectively) were collected from each forest type in October 2014. Both litter types were brought back to the laboratory and dried at 50°C for 48 hours. After drying, the litters were carefully sorted to remove other organic matter (e.g. twigs) and then stored at room temperature in paper bags prior to litter bag construction. Approximately 2 g of oven-dried pine or oak litter was weighed and placed in litter bags constructed of polyurethane 2 mm mesh (Industrial Netting, Minneapolis, MN, USA, Product #XN3234) *ca.* 12 x 12 cm in dimension and heat sealed closed. For the first experiment, litter bags matching the canopy composition of the forest (i.e. pine litter in the pine site and oak litter in the oak site) were incubated at the soil-litter layer interface in each plot, with the incubation starting on July 15, 2015. For the second experiment, both pine and oak litter (in separate litter bags) were incubated in the same plots, with the incubation commencing on July 15, 2016. In each experiment, the litter bags were incubated in each plot for 2, 4 and 12 months. Upon harvesting, the litter bags were placed in sterile plastic bags and transported back to the laboratory where the litter was re-dried at 50°C until the mass was stable. The remaining mass of the decomposed litter was determined using plastic weighing trays sterilized with 70% ethanol. The dried litter was then stored in labeled sterile plastic bags at −20°C ahead of elemental and molecular analyses.

### Litter chemistry

Lignin, cellulose, and hemicellulose concentrations were measured for initial and 12-month incubated litter using an ANKOM Fiber Analyzer (Ankom Technology, Macedon, New York, USA: Hobbie 2008). The C and N content for samples incubated for 12 months was assessed via dry combustion (Costech ECS 4010 Elemental Analyzer, Valencia, CA, USA) at the University of Minnesota. Elemental contents of the mass remaining were then calculated by multiplying mass by concentration.

### Fine root in-growth

To determine the effectiveness of the trenching treatment in terms of reduction of fine root ingrowth, we placed in-growth cores (5 cm x 15 cm) containing sieved soil just inside and outside the edge of the trenched plots in each block and incubated them for 30 days in July 2016. The in-growth cores were brought back to the lab and the soil was sieved with a 2 mm sieve to collect total root biomass. Root biomass was then rinsed in water and fine roots were identified as herbaceous or woody. Only fine woody roots, which dominated the root pool, were dried, weighed and included in the analysis.

### Soil nutrient availability

To assess soil inorganic nutrient availability in the untrenched and trenched plots, we incubated 3 pairs of plant root simulator (PRS) probes (Western Ag Innovations; Saskatoon, SK, Canada) in each plot from June 2 – July 6, 2017, which corresponds to a period of high plant productivity at Cedar Creek. The cation and anion PRS probes were oriented vertically in the top 10 cm of the soil and across the soil plots. After the incubation the PRS probes in each plot were pooled and sent to Western Ag Innovations for processing.

### Fungal community identification

Genomic DNA was extracted from all litter samples using MoBio PowerSoil kits (MoBio, Carlsbad, CA, USA). Prior to extraction, a ca. 100 mg subsample was homogenized via bead beating in 2 ml tubes containing 3 one mm zirconia/silica beads (BioSpec Products, Bartlesville, OK, USA). In addition, genomic DNA from soil cores collected from untrenched and trenched plots 12 months after establishment were also extracted using the same extraction method. To characterize fungal community composition, rDNA of the ITS1 region was PCR amplified using a barcoded fungal-specific ITS1F-ITS2 primer set, following the reagent and cycling conditions detailed in Smith and Peay (2014). While this primer set has been critiqued for not amplifying members of the genus *Mycena* (Lindahl and Tedersoo, 2016), we found many sequences that could be successfully matched to this genus, so do not think this primer choice resulted in significant methodological bias. A 25 fungal species mock community detailed in Nguyen et al. (2015) was also included. Amplified products were cleaned and normalized individually using Charm ‘Just-a-Plate’ kits (Charm, San Diego, CA, USA). The samples from each experiment were pooled into individual libraries and sequenced at the University of Minnesota Genomics Center using 250 bp paired-end V2 MiSeq Illumina chemistry (Illumina, San Diego, CA, USA).

Fungal sequences were processed using the amptk pipeline version 1.1 (Palmer et al., 2018). Briefly, the forward and reverse sequences in each sample were demultiplexed, the primers removed, and then denoised using the UNOISE3 algorithm (Edgar, 2016). The resulting ‘inferred sequences’ (a.k.a. exact sequence variants) were clustered into operational taxonomic units (OTUs) at 97% similarity using VSEARCH (Rognes 2016). Taxonomy was assigned using a ‘last common ancestor’ approach of global USEARCH, UTAX, and SINTAX alignments against the UNITE v7.2.2 database (Kõljalg et al., 2013). To remove possible sequences caused by index bleed, a 0.5% filtering was applied to each sample. In addition, for each OTU, any sequence reads present in the PCR negative controls were subtracted from read abundances present in the litter samples (Nguyen et al., 2015). Finally, the mock community was also used to determine the level at which unexpected sequence reads were encountered resulting in all cells, with values less than 4 sequence reads being zeroed.

Following Sterkenberg et al. (2015), we assigned all OTUs belonging to the Eurotiales, Hypocreales, Morteriellales, Mucorales, Saccharomycetales, Tremellales and Sporidiales as ‘Molds & Yeasts’ to better reflect the r-selected life history strategies of these groups and distinguish them from soil and litter associated saprotrophic fungi. The remaining OTUs were assigned to saprotrophic, ectomycorrhizal, other symbiotrophic (e.g. arbuscular and ericoid mycorrhizal fungi), and pathotrophic guilds modes using FUNGuild (Nguyen et al. 2016). When possible, the top 50 most abundant unassigned OTUs (due to missing genus taxonomy) were assigned manually to either EM or saprotrophic guilds using criteria detailed in Fernandez et al. (2017). A list of guild assignments are provided in Table S3. From the 4,294,076 sequence reads passing the quality filtering steps, 2,423,625 sequence reads (mean = 55% of reads/sample) could be assigned to functional guilds. Because of variation in sequence read depth per sample, the final EM and saprotroph dataset was normalized to proportional abundances or Hellinger transformed (sqrt relative abundance). Raw sequence read files are available in NCBI SRA accession: [released upon manuscript acceptance].

### Statistical analyses

The effect of site, trenching and potential interactions on root in-growth rates, soil moisture, soil fungal guild abundance, soil nutrient availability, and incubated litter chemistry (12 months) were assessed using linear mixed models with block nested in site as a random factor. For experiment 1, the effect of trenching and incubation time on mass remaining was tested with linear mixed models for each litter type with block nested in site as a random effect. For experiment 2, the effects of site, trenching, and incubation time on litter mass remaining were tested using linear mixed models for each of litter type with block nested within site as a random effect. All mixed models were run using the ‘lme’ function in nlme package in R. Soil nutrient availability data and all litter mass remaining data was ln transformed prior to analysis to satisfy the linearity assumptions.

To visualize HTS fungal community data we used ‘amp_ordinate’ functions in the *ampvis2* package in R to construct non-multi dimensional scaling plots. The effects of trenching and incubation time on fungal community composition were assessed using factorial permutational analyses of variance (PERMANOVA). Prior to each PERMANOVA, data were Hellinger-transformed and pair-wise distances were calculated based on Bray-Curtis dissimilarity. To further assess whether significant results detected by the PERMANOVA analyses were due to shifts in composition or heterogeneity, betadisper tests were run on significant predictor variables. Finally, ANCOVAs were used to detect the effect of the trenching treatment on the C and lignin content per unit N for each of the litter types after 12 months of incubation.

## Results

*Soil* Soil fungal community composition differed significantly among the two sites (Fig. 1a, PERMANOVA; Site: *F*1,42 = 12.43; *P* < 0.0001), which was not caused by differences in community heterogeneity (betadisper: Site: *P* = 0.67). In both the pine and oak sites, the soil fungal communities were dominated by EM fungal taxa (Fig. 2). The most abundant EM genera in the pine site were *Tomentella*, *Russula*, and *Inocybe* (Fig. 1b), whereas *Russula*, *Amanita*, *Scleroderma*, *Tomentella*, *Cortinarius* and *Cenococcum* were the dominant EM fungal genera in the oak site (Fig. 1c). The non-mycorrhizal (i.e. saprotrophic, molds & yeasts, pathotrophic) fungal genera dominating the pine site soil community included *Lepiota*, *Chalara*, *Trechispora* and *Mortierella,* while *Mortierella, Cladophialophora,* and *Vararia* dominated the oak site soil communities (Fig. 1c). Trenching had a marginal effect on fungal community composition (PERMANOVA; Trenching: *F*1,42 = 1.37, *P* = 0.108), but significantly reduced the abundance of EM fungi in soils in both sites (Trenching: *F*1,30 = 11.37, *P* = 0.002; Fig. 2). In contrast, trenching increased the abundance of molds & yeasts in both sites (Trenching: *F*1,30 = 18.70, *P* = 0.0002), and saprotrophic fungi in the pine site but not the oak site (Site × Trenching: *F*1,30 = 6.05, *P* = 0.02; Fig. 2).

**Figure 1.**
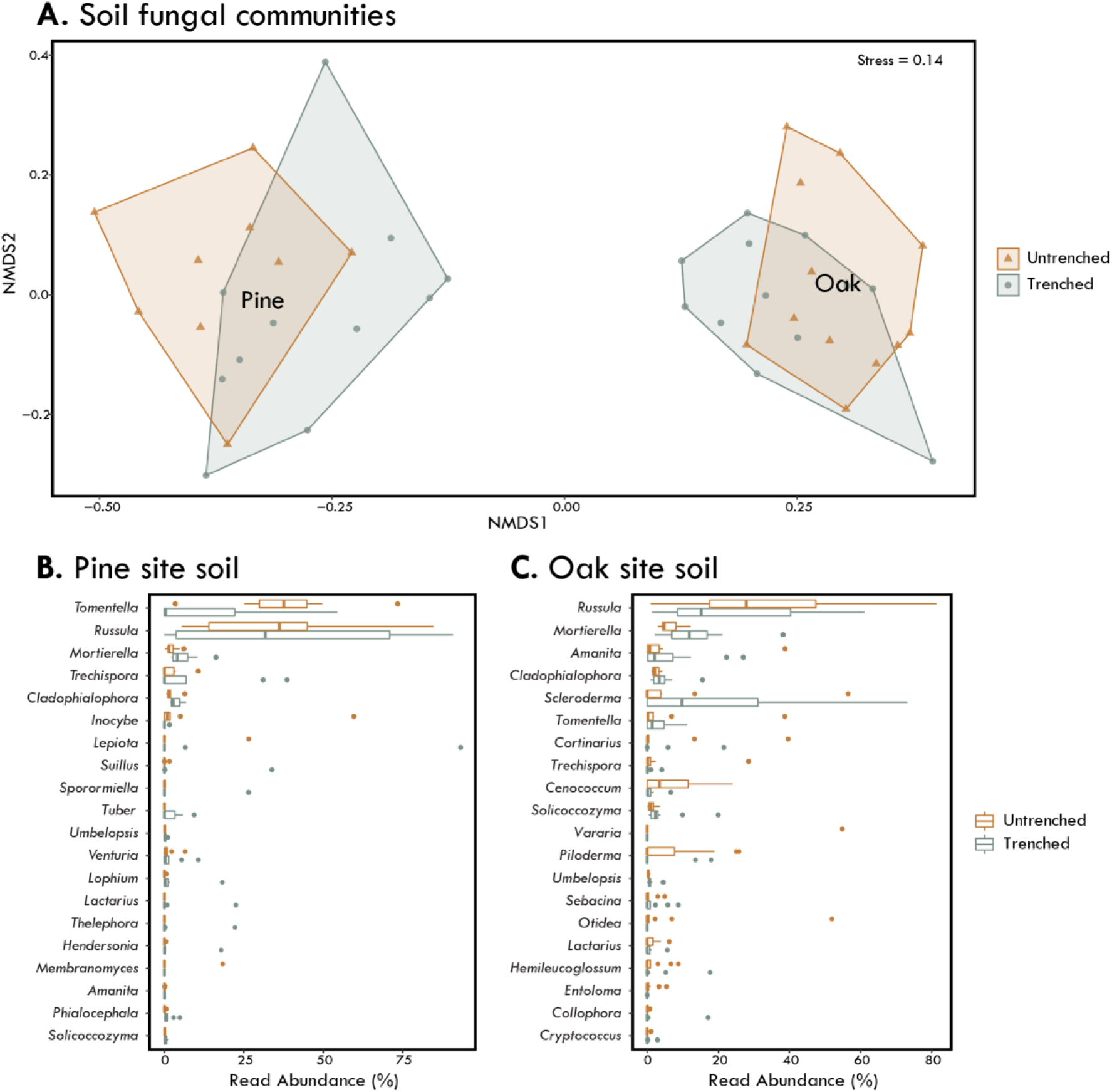
Soil fungal communities at the oak and pine sites and their response to trenching. Non-metric multidimensional scaling plots of soil fungal communities present in the oak and pine sites. Each site label represents the site centroid **(A).** Samples and frames are colored by untrenched (orange) and trenched (gray) treatments to help visualize compositional differences. Boxplots of relative read abundance of the top 20 most abundant fungal genera present in the in untrenched (orange) and trenched (gray) treatments in the pine site soils **(B)** and the oak site soils **(C)**. (n=44)

**Figure 2.**
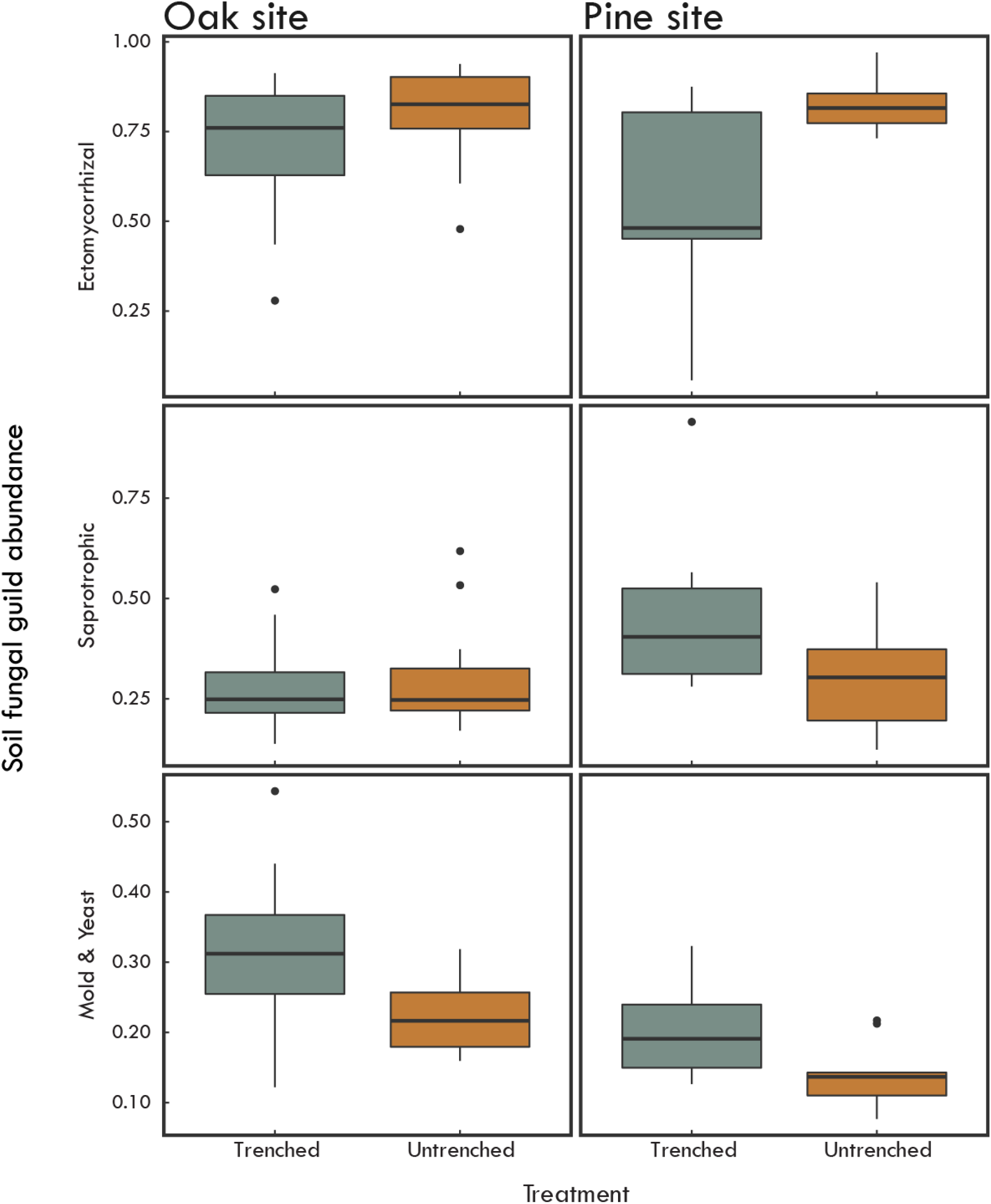
Boxplots of ectomycorrhizal, saprotrophic, and mold & yeasts abundance in the untrenched (orange) and trenched (gray) treatments after 12 months oak and pine site soils. (n=44).

Root in-growth rates were approximately twice as fast in the oak site compared to the pine site, based on measurements in the untrenched plots (Fig. S2). Trenching significantly reduced root in-growth frequency and decreased mean in-growth rate in both sites (Trenching: *F*1,10 = 8.67; *P* = 0.014; Fig. S2). There were no significant differences between the O-layer (Site: *F*1,9 = 0.04; *P* = 0.84) and mineral soil moisture (Site: *F*= 1.73, *P*= 0.24) between the two sites. Trenching increased mineral soil moisture where from 3.6% to 5.9% water on average (Trenching: *F* = 22.17; *P* = 0.001). Although trenching significantly reduced root in-growth rates and mineral soil moisture at both sites, there was no effect of this treatment on O-layer moisture content in either site during the growing season (Trenching: *F* = 0.0436, *P* = 0.8373). EM root in-growth rate was positively correlated with EM fungal abundance in both sites (Fig S1). Conversely, EM root in-growth rate was negatively correlated with saprotrophic fungal abundance in the pine site but no relationship was detected in the oak site (Fig S1). With respect to inorganic N availability, the trenching significantly increased soil N availability in the pine site but had no effect in the oak site (Site × Trenching: *F*1,10 = 6.48; *P* = 0.029, Fig. 3). Conversely, phosphorous availability was significantly higher in the untrenched plots in both sites (Trenching: *F*1,10 = 6.75; *P* = 0.026, Fig. 3).

**Figure 3.**
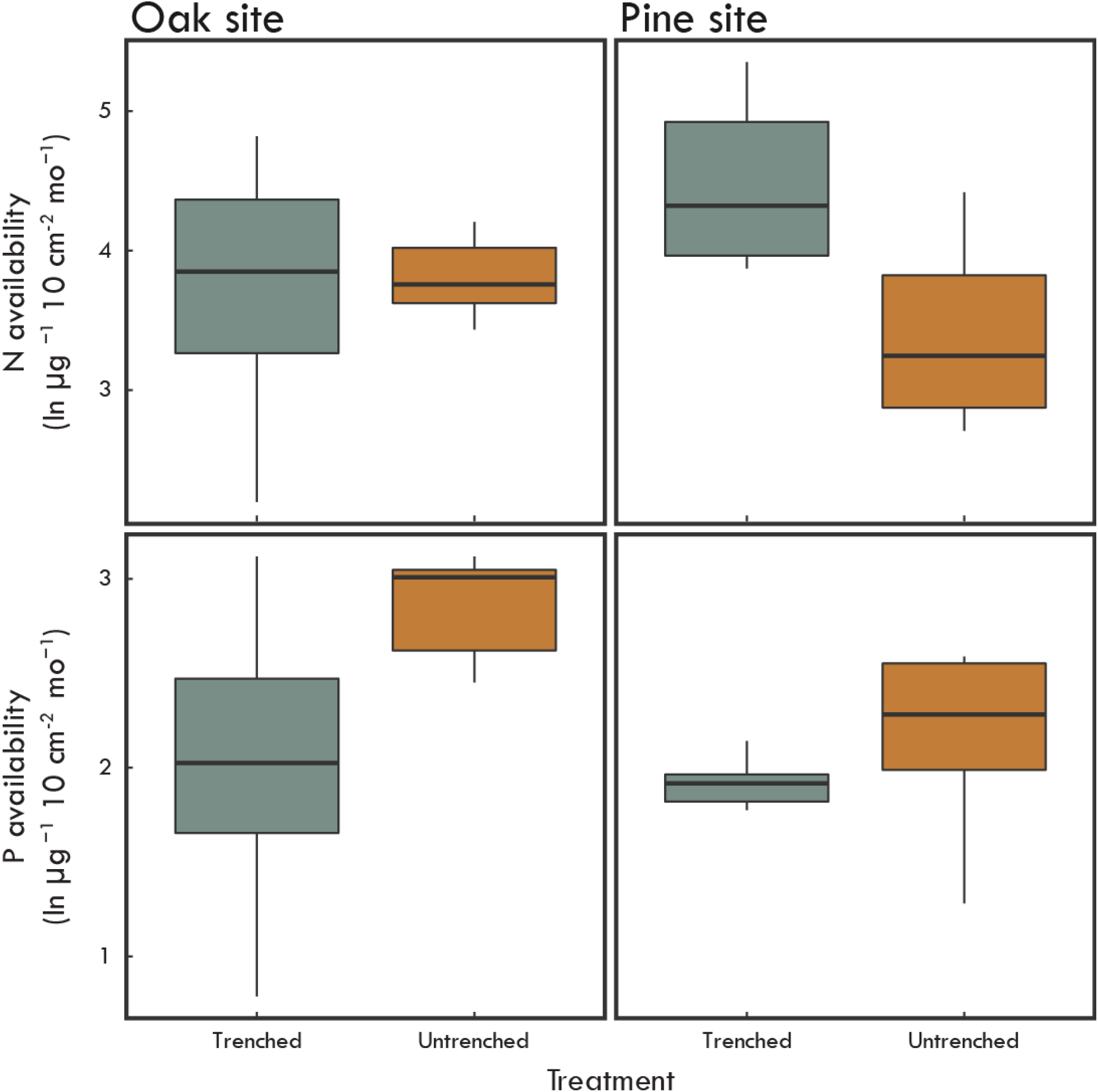
Soil nitrogen and phosphorus availability in the untrenched (orange) and trenched (gray) treatments after 12 months oak and pine sites soils. (n=24)

EM root in-growth rates were marginally negatively correlated with soil N availability in the pine site (*P* = 0.10; R^2^ = 0.26), but no significant relationship was detected in the oak site (Fig 4a). Similarly, the ratio of ectomycorrhizal-to-saprotrophic fungal abundance was negatively correlated with soil N availability in the pine site (*P* = 0.008; R^2^ = 0.56), but again no significant relationship was detected in the oak site (Fig. 4b).

**Figure 4.**
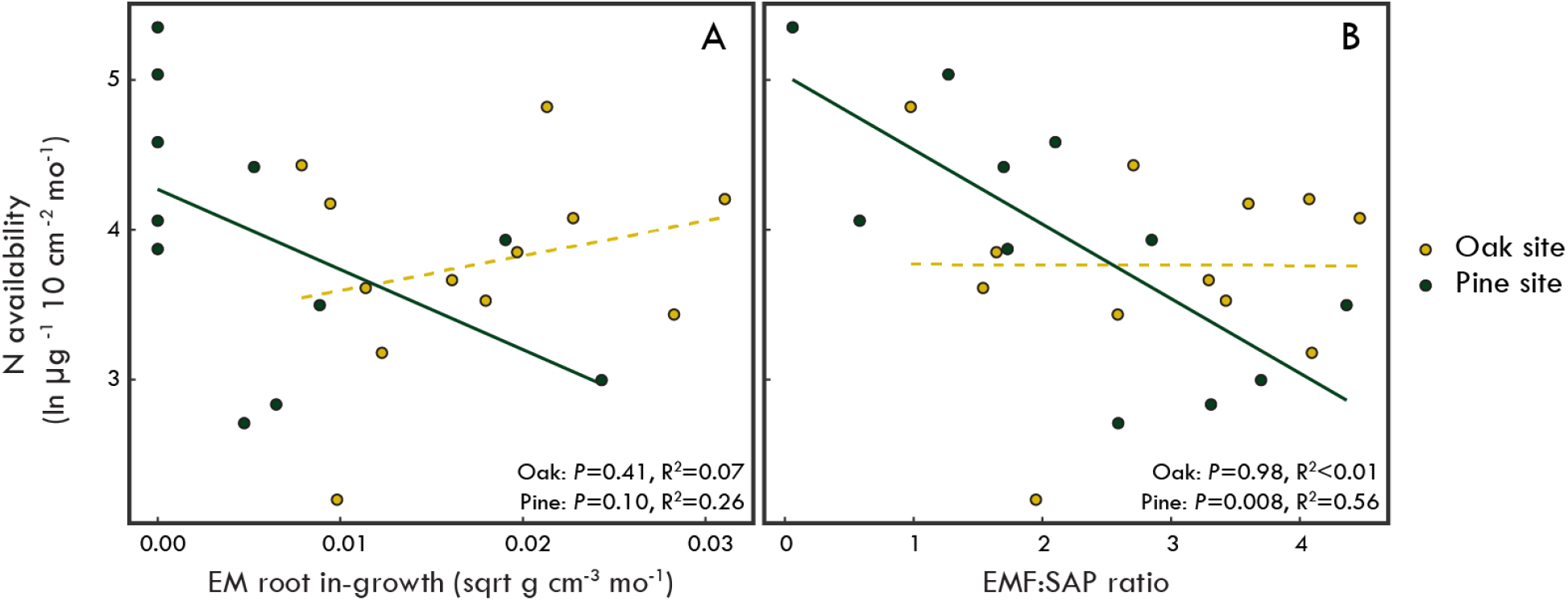
Relationship between soil nitrogen availability and ectomycorrhizal root in-growth rate (A) and soil ectomycorrhizal-to-saprotrophic abundance ratio (EMF:SAP) (B). Data points and regression lines are colored by oak (yellow) and pine (green) site soils. Both untrenched and trenched treatments were included in the analyses. (n=23)

### Litter decomposition

#### Experiment 1

There was no effect of trenching on oak litter mass loss rates (Table 2), in fact, litter mass loss rates in the trenched oak site plots were actually lower at the 2 month time point (Fig. S3), suggesting that, at least initially, roots and/or EM fungi may be involved in accelerating litter decomposition rather than suppressing it in this site. Conversely, the trenching treatment had an increasingly positive effect on decomposition rates of pine litter incubated in the pine site (Table 2; Fig. S3).

**Table 1.**
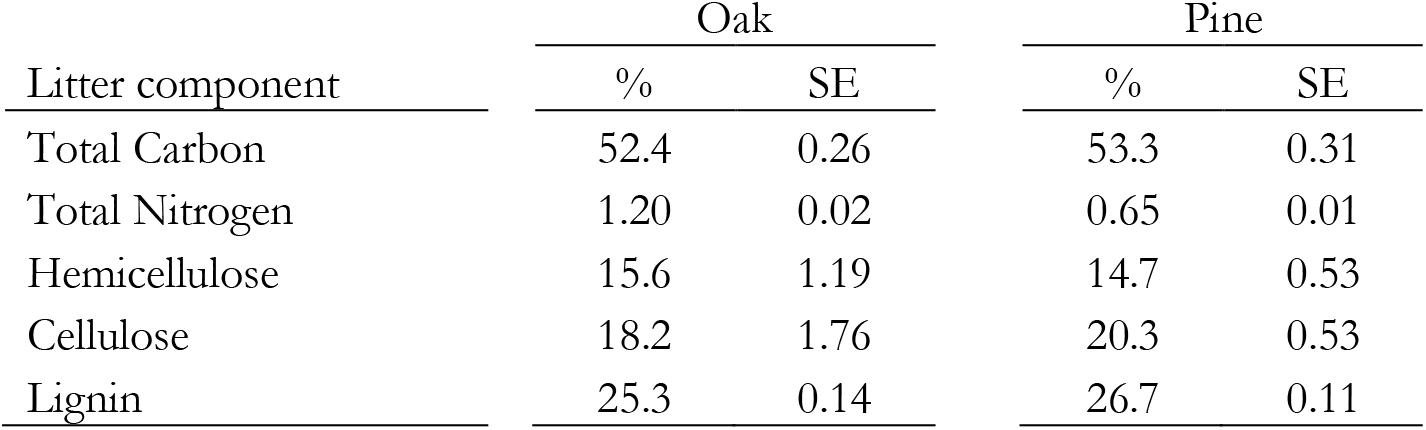
Initial litter chemistry.

**Table 2.**
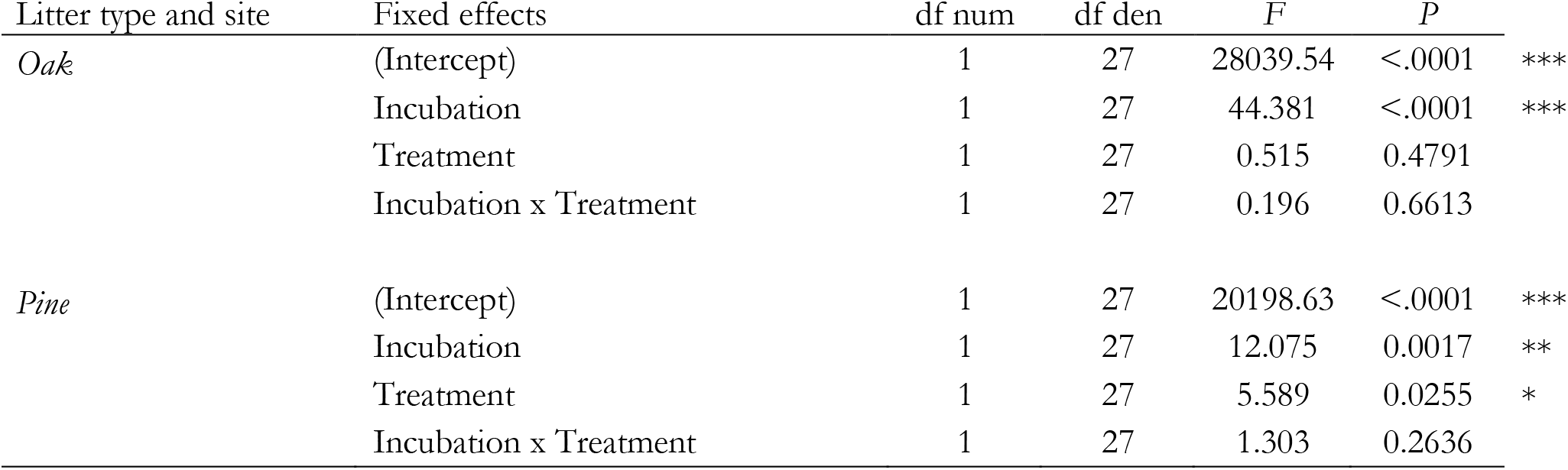
Experiment 1 effects tests from mixed models explaining litter mass remaining.

#### Experiment 2

The effect of trenching on the mass loss was different for the two litter types and also depended on the site it was incubated in (Table 3). Trenching significantly increased pine litter mass loss rates when incubated in the pine site after 12 months (Fig. 5d), but not when pine litter was incubated in the oak site (Fig 5c). In contrast, oak litter mass loss rates were not significantly affected by trenching in either site (Table 3; Fig. 5a,b). Oak litter, which had lower lignin and higher N concentrations (Table 1), had, on average, higher rates of mass loss compared to pine litter, after 12 months of decomposition. However, litter mass loss rates were, on average, higher in the pine site relative to the oak site.

**Table 3.**
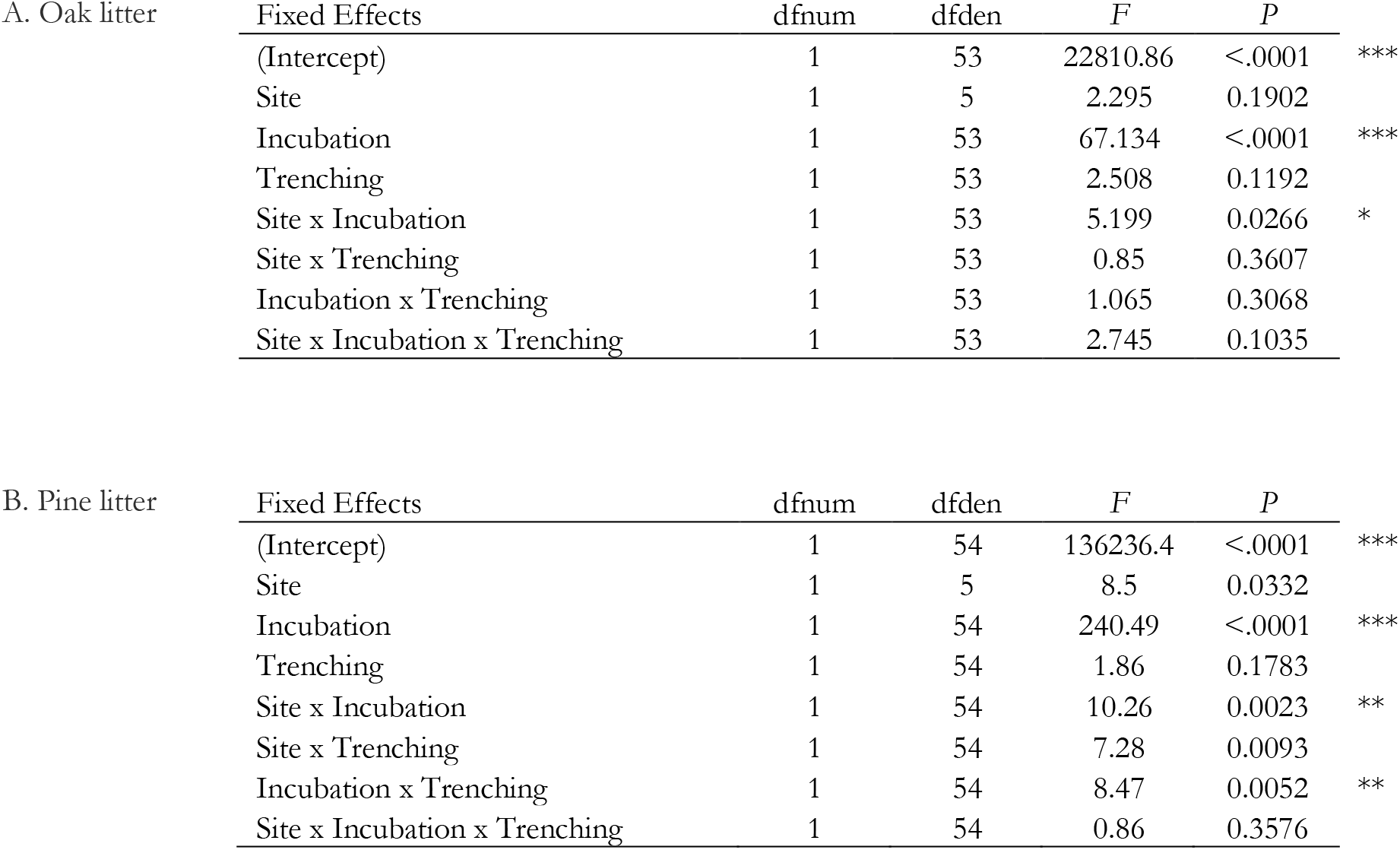
Experiment 2 effects tests from mixed models explaining oak (A) and pine (B) litter mass remaining.

| A. Oak litter | Fixed Effects | dfnum | dfden | *F* | *P* |  |
| --- | --- | --- | --- | --- | --- | --- |
|  | (Intercept) | 1 | 53 | 22810.86 | <.0001 | *** |
|  | Site | 1 | 5 | 2.295 | 0.1902 |  |
|  | Incubation | 1 | 53 | 67.134 | <.0001 | *** |
|  | Trenching | 1 | 53 | 2.508 | 0.1192 |  |
|  | Site x Incubation | 1 | 53 | 5.199 | 0.0266 | * |
|  | Site x Trenching | 1 | 53 | 0.85 | 0.3607 |  |
|  | Incubation x Trenching | 1 | 53 | 1.065 | 0.3068 |  |
|  | Site x Incubation x Trenching | 1 | 53 | 2.745 | 0.1035 |  |
| B. Pine litter | Fixed Effects | dfnum | dfden | *F* | *P* |  |
|  | (Intercept) | 1 | 54 | 136236.4 | <.0001 | *** |
|  | Site | 1 | 5 | 8.5 | 0.0332 |  |
|  | Incubation | 1 | 54 | 240.49 | <.0001 | *** |
|  | Trenching | 1 | 54 | 1.86 | 0.1783 |  |
|  | Site x Incubation | 1 | 54 | 10.26 | 0.0023 | ** |
|  | Site x Trenching | 1 | 54 | 7.28 | 0.0093 |  |
|  | Incubation x Trenching | 1 | 54 | 8.47 | 0.0052 | ** |
|  | Site x Incubation x Trenching | 1 | 54 | 0.86 | 0.3576 |  |

**Figure 5.**
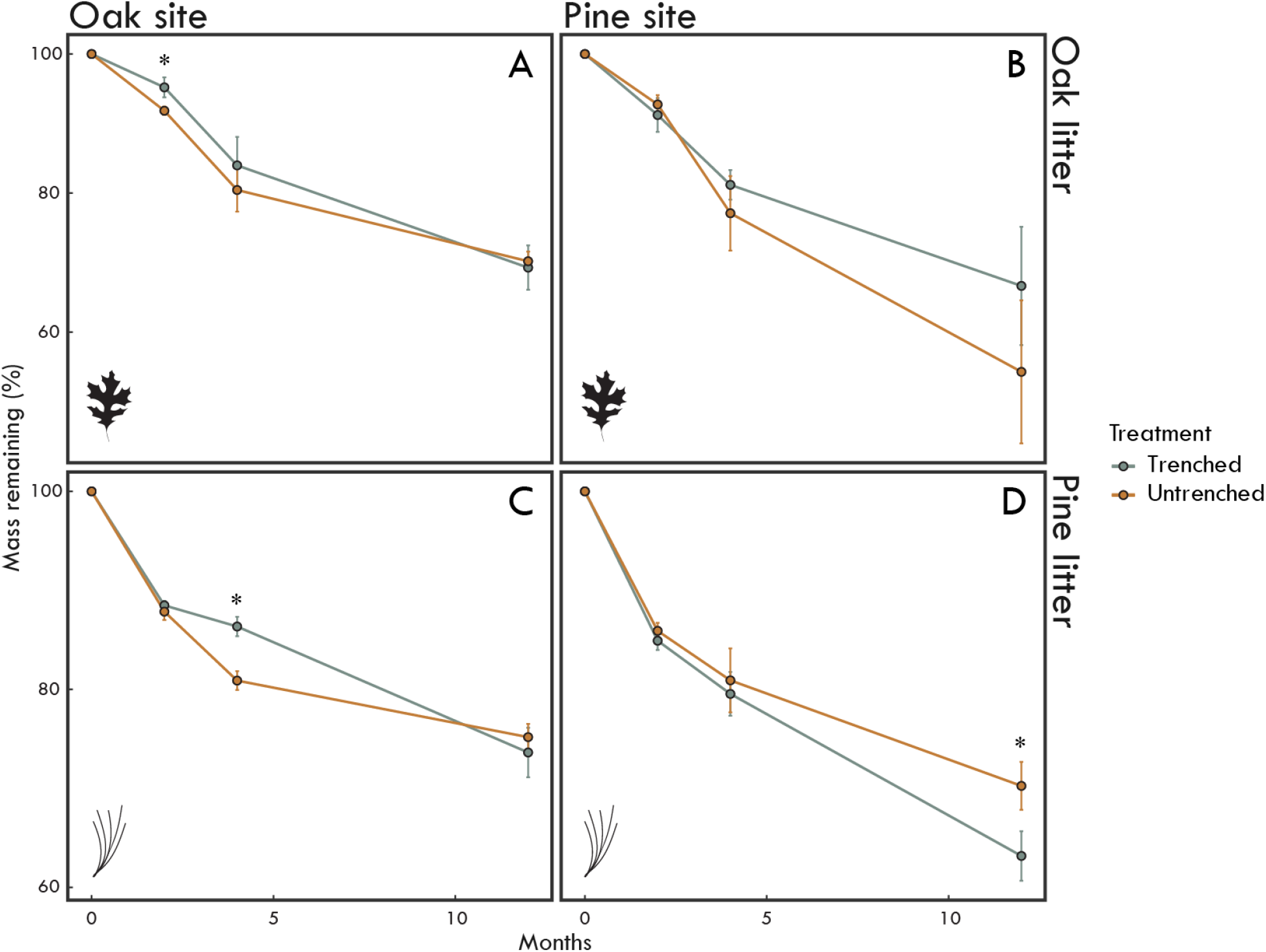
Percent mass remaining (Mean ± SE) of oak **(A. & B.)** and pine **(C. & D.)** litter incubated in the untrenched (orange) and trenched (gray) treatments for 2, 4 and 12 months in the oak **(A. & C.)** and pine site **(B. & D.)** (n=196)

The N concentration of the incubated litter differed significantly depending on litter type and the site in which it was incubated (Site × Litter: *F*1,28 = 7.63, *P* = 0.010). This interaction was driven by oak litter, which had a higher initial N concentration compared to pine litter (Table 1), but had significantly lower post-incubation N concentrations than pine litter when decomposed in the pine site regardless of trenching (Table S4; Fig. S6). There were no significant effects of trenching on litter N concentration (Table S4), but the C:N ratios of the incubated litters were higher in the untrenched compared to trenched plots (Trenching: *F*1,28 = 4.54; *P* = 0.042). In addition, there was an interaction between litter type and site; each of the litters had slightly higher C:N ratios when incubated in the site with non-matching canopy composition (Site × Litter: *F*1,28 = 9.31; *P* = 0.005).

While the C concentration of the incubated litters largely had a consistent response to trenching within sites, further fractionation of the C remaining after incubation revealed important differences in the recalcitrant lignin fraction in response to trenching among the litter types and sites (Site × Litter × Trenching: *F*1,28 = 5.14; *P* = 0.031). Pine litter always had higher lignin concentrations in untrenched plots compared to trenched plots no matter which forest type in which it was incubated (Fig. S6). Conversely, oak litter incubated in the oak site had significantly lower lignin concentration in the untrenched plots compared to trenched plots, but when incubated in the pine site the opposite was observed (Fig. S6). Lignin loss during the incubation was also affected by a litter by treatment interaction (*P* < 0.001), with pine litter losing 14 and 15% *more* lignin in the trenched than untrenched plots in the oak and pine sites, respectively. Conversely, oak litter lost 15 and 4 % *less* lignin in the trenched than untrenched plots in the oak and pine sites, respectively. Finally, litter types had notably different responses to trenching in terms of remaining C content per unit N (Table S5; ANCOVA: N Content × Litter × Trenching: *F*1,20 = 5.62; *P* = 0.003) as well as remaining lignin content per unit N (Table S5; ANCOVA: N Content × Litter × Trenching: *F*1,20 = 12.20; *P* = 0.002). Pine litter had higher C and lignin content per unit N in the untrenched than trenched plots (Fig. 6b,d), whereas for oak litter there was no difference in terms of C content per unit N by treatment and lower lignin content per unit N in the untrenched than trenched plots (Fig. 6a,c).

**Figure 6.**
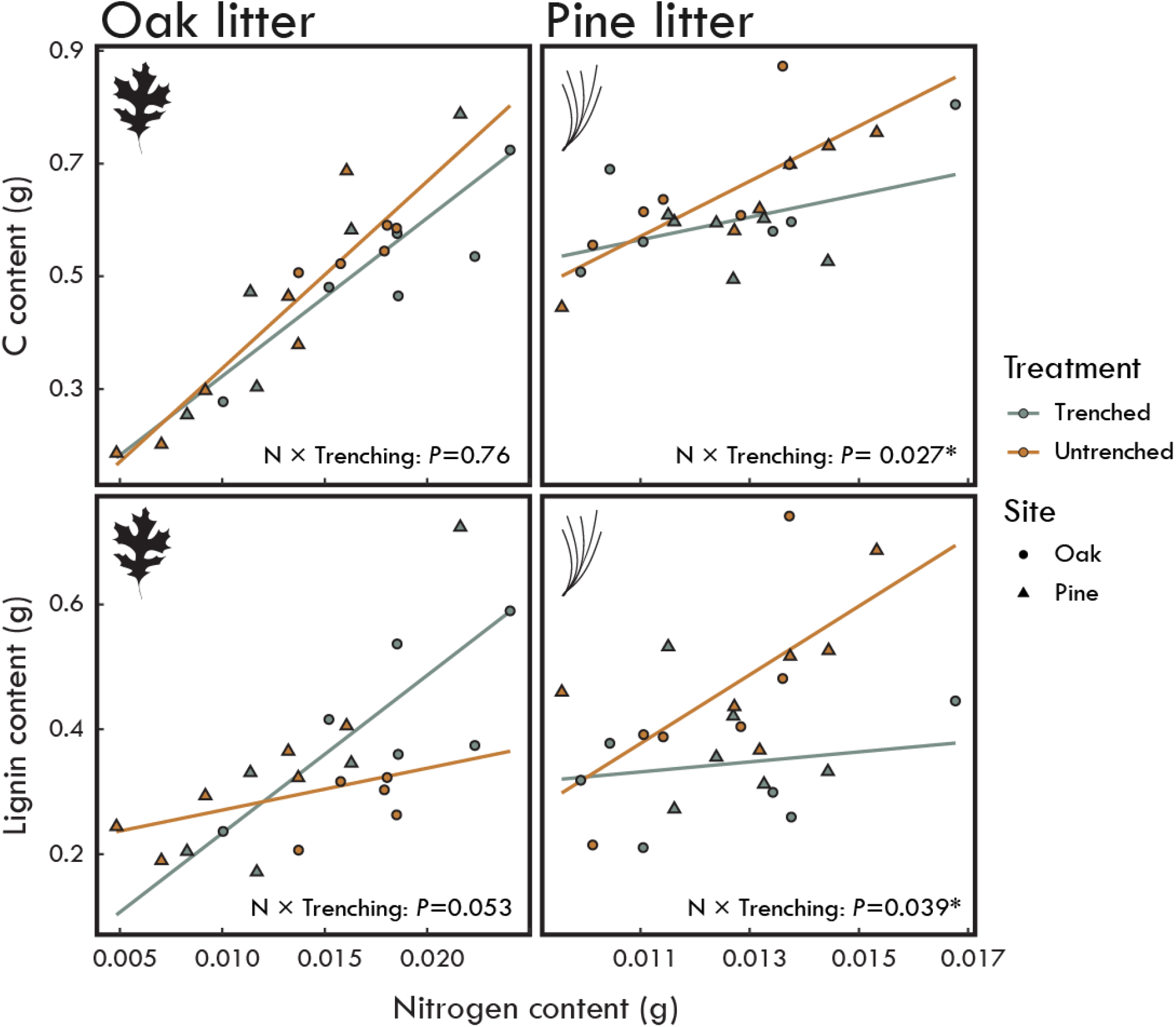
The relationship between oak and pine litter carbon content and lignin content with nitrogen content by untrenched (orange) and trenched (gray) treatments. Differences in slopes among the treatments were determined with ANCOVA models. (Oak litter n=21; Pine litter: n=23).

Litter and forest types were also important determinants of fungal guild abundances over the course of the incubation (Table S6; Fig. S4). Oak litter incubated in both forests was dominated by saprotrophic fungi, although the change in their relative abundance over the incubation varied somewhat between forest types. The abundances of all other fungal guilds associated with the oak litter, including EM fungi, were very low compared to saprotrophic fungi (Fig. S4). Conversely, abundances of various fungal guilds colonizing the pine litter were far more even in both sites and pine litter had notably higher EM fungal colonization in the untrenched treatment of the pine site (Fig. S4).

The specific taxonomic composition of the fungal communities colonizing litter was dependent on site, litter type, and incubation time (PERMANOVA: Site × Litter type × Incubation: *F* = 1.93; *P* = 0.035), with trenching again having marginal effects (PERMANOVA: Trenching: *F* = 1.606; *P* = 0.07). Notably, members of the EM genus *Tomentella* were abundant in the pine litter incubated in the untrenched plots in the pine site and were dramatically reduced in the trenched plots (Fig. S5). When the oak litter was incubated in the pine site *Tomentella* was also somewhat abundant after 12 months of incubation, but nowhere near the levels seen in the pine litter (Fig S5). Conversely, when the pine litter was incubated in the oak site, the EM genus *Amanita* was moderately abundant in the untrenched plots yet practically absent in the trenched plots. Many of the dominant non-mycorrhizal fungi showed litter-specific associations and abundance patterns (e.g. *Talaromyces* and *Pezicula* for oak litter, *Lophium* and *Xenopolyscylalum* for pine litter (Fig. S5), but generally had similar or higher abundances in the trenched relative to the untrenched plots.

## Discussion

### Generality and context dependency of the ‘Gadgil effect’

While the results from this study support earlier findings that EM fungi can suppress leaf litter decomposition rates, they also question the generality of this phenomenon. In soils at the pine site, we found a clear and consistent negative relationship between EM fungi and N availability. Specifically, when EM fungal in-growth and abundance were reduced by trenching, there was significantly faster litter decomposition, which is consistent with the ‘Gadgil effect’. Conversely, in soils at the oak site, we found neutral relationships between EM fungi and soil N availability, and trenching had no effect on litter decomposition rates. These results, while based at the local scale, parallel inconsistencies observed at larger spatial scales in the literature (Fernandez & Kennedy, 2016). Some experiments have shown strong negative effects of EM fungi on OM decomposition (Gadgil & Gadgil, 1971; Gadgil & Gadgil, 1975; Berg & Lindberger, 1980; Fisher & Gosz, 1986; Koide & Wu, 2003; Moore et al., 2015; Averill & Hawkes, 2016; Sterkenberg et al. 2017; Maaroufi et al., 2019), while others have reported no effects (Harmer & Anderson, 1985; Staaf, 1988; Mayor & Henkel, 2006; McGuire et al., 2010; and Brzostek et al., 2015), or even positive effects (Zhu & Ehrenfeld, 1996). Comparing results across those studies is complicated by the fact that they are conducted in different ecosystems with many covariables (e.g. climate and soil properties), which may interact with the effect of EM fungi on soil biogeochemical cycling. Here, by working in sites in close physical proximity, we held climate and edaphic factors functionally constant, thereby isolating the effects of tree hosts and EM fungal communities on forest C and N cycling. The strong differential responses we observed here among sites differing in both plant and fungal community composition point to the importance of understanding the biotic underpinnings of fungal interguild competition and the ‘Gadgil effect’.

### Litter-Fungal community interactions

Teasing apart the interactions between top-down (litter chemistry) and bottom-up (EM fungi) controls over litter decomposition is necessary for a complete understanding of the context dependency of the ‘Gadgil effect’. Fernandez & Kennedy (2016) hypothesized that N availability and narrow C:N ratio of litter inputs may favor higher carbon use efficiency among free-living saprotrophs and reduce the effectiveness of EM fungi in acquiring organic N. This was supported by results from Kyaschenko et al., (2017), who found correlative evidence along a fertility gradient that competitive interactions between EM and saprotrophic fungal guilds may be partially driven by N availability. Recently, Smith & Wan (2019) modelled the competitive interactions between EM and saprotrophic fungi and their consequences on soil C and N cycling by applying resource ratio theory (Tillman 1982) to better understand and predict context dependencies of the ‘Gadgil effect’. The model predicted that litter decay rates were decelerated only when the starting substrate was energetically unfavorable to saprotrophic fungi (e.g. wide C:N ratio, high Lignin:N ratio). While our results support this predicted top-down control of fungal competitive interactions, they also suggest that EM fungal community composition and inherent functional differences (e.g. enzymatic suites, exploration strategy, vertical niche preference) are probably of equal importance. Specifically, we found that the higher quality oak litter was always dominated by saprotrophs, had low EM fungal colonization and decay rates that were largely unresponsive to the trenching treatment. Additionally, only the lower quality pine litter (wider initial C:N and Lignin:N ratios), when incubated in the pine site was dominated by EM fungi and led slowed decay rates. Conversely, when pine litter was incubated in the oak site, we observed an opposite trend. Because the two sites differed significantly in fungal community composition, this suggests that there are important functional differences among members of these EM fungal communities. Finally, over the 12 month incubation we observed widening of these ratios in the pine litter in the untrenched plots compared to the trenched plots, suggesting that EM fungi were involved in decoupling of C and N cycling of the litter, which was not observed in the oak litter. Taken together, these findings support a view that EM-mediated deceleration of leaf litter is likely dependent on both substrate chemistry and EM fungal community composition (Figure 7).

**Figure 7.**
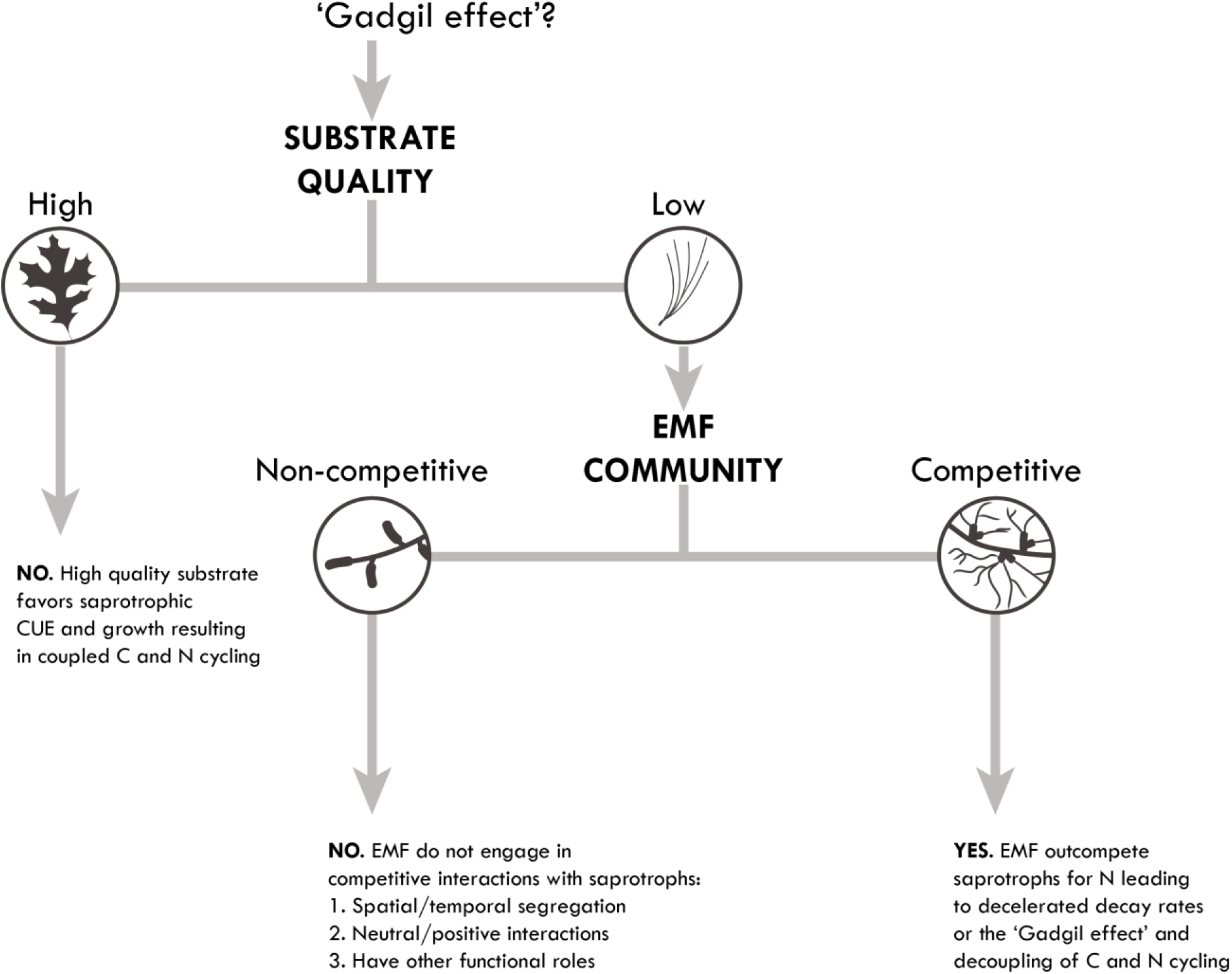
Conceptual summary of the effects of litter substrate quality and EM fungal community composition on the presence or absence of the ‘Gadgil effect’.

### Functional diversity of ectomycorrhizal fungi

Despite the litter/humus layers of soils being generally thought of as the exclusive domain of saprotrophic fungi (Lindahl et al., 2007), a growing number of studies have demonstrated that EM fungi frequently colonize these substrates as well (Dickie et al., 2002; Genney et al., 2006; Hobbie, et al., 2014; Anderson et al., 2014). Given the functional diversity that exists within and across EM fungal communities (Read & Perez-Moreno, 2003; Finlay 2008; Koide et al., 2007; Koide et al., 2014), it is not particularly surprising that the litter decomposition patterns we observed imply important functional differences among the EM fungi associated with each forest. Interestingly, however, unlike previous work focusing on the genus *Cortinarius* (Clemmensen et al. 2015, Kyashenko et al. 2017), the suppression of free-living fungi and litter decay rates was consistently associated with increased abundance of EM OTUs in the genus *Tomentella*. Although this genus has not been previously recognized as one with high SOM degradation potential, a number of studies indicate that EM root tips colonized by these fungi are capable of producing a wide range of extracellular enzymes used to breakdown proteins, polysaccharides, and organic forms of P (Courty et al., 2005; Tedersoo et al., 2012). In addition, Pena et al., (2013) demonstrated that *Tomentella badia* colonized ^15^N-labeled beech leaf litter (*Fagus sylvatica*) and the enrichment of associated EM roots were approximately four times higher than any of the other EM fungi examined. That result indicates that at least some members of this genus are highly capable of mobilizing organic N from leaf litter. A second intriguing finding with regard to *Tomentella* litter colonization was its very limited colonization of the oak litter incubated in the pine site. Again, we suspect this may in part be due to the oak litter having relatively labile chemistry that favored free-living fungi (Smith & Wan 2019). We further hypothesize that the EM fungal community in our oak site may therefore be comprised of EM fungi that favor acquisition of mineral bound N and/or priming of N mineralization rates via C exudation by roots and EM fungi (Phillips et al., 2012). Finally, it is important to emphasize that while *Tomentella* species appear to be driving a ‘Gadgil effect’ in this system, other EM fungal taxa may have similar effects and influence on C and N cycling in other systems. We therefore recommend further investigation into EM fungal community composition and linkages with the ‘Gadgil effect’ in other ecosystems with the hope of identifying common functional traits influencing the phenomenon.

While we believe our results provide important insights regarding the role of EM fungi in mediating leaf litter decomposition, a number of caveats should be noted. In our study system (Cedar Creek Ecosystem Science Reserve), the soils lack a well-developed organic layer, which may increase the shared realized niche space between EM fungal and saprotrophic guilds and thereby intensifying interactions (Bödeker et al., 2016). As noted above, this possibility was supported by the recent findings of Kyaschenko et al., (2017), who showed strong correlative evidence that competition between EM and saprotrophic guilds may be partially driven by the degree of organic layer development. Like other recent studies of the ‘Gadgil effect’ (Averill & Hawkes, 2016; Sterkenberg et al., 2018), our inferences regarding EM and saprotrophic fungal abundance are based on sequence read counts. While we clearly acknowledge the semi-quantitative nature of this metric (Amend et al., 2010), we believe the consistent and sizable declines in EM-to-saprotrophic fungal ratios in the trenched plots and the positive correlation between EM root in-growth indicates the differences in C and N cycling we observed were related to significant changes in EM abundance. Additionally, while the results we observed are consistent with the broader literature noting the preferential presence of a ‘Gadgil effect’ in conifer forests, further investigation of the phenomenon at larger spatial scales is needed to confirm this pattern. Finally, we recognize that the suppression of litter mass loss we observed does not necessarily translate directly into greater C stocks in soil (Schmidt et al., 2011). For instance, while the suppression of litter and particulate organic matter decay rates by EM fungi may lead to increased C stocks in those SOM fractions, they may ultimately lead to a slower accrual of *total* C stocks due to reductions in the rate of mineral associated organic matter (MAOM) formation. This slowing would be driven by declines in microbial CUE, biomass production, and stabilization necromass C to mineral exchange sites (Cotrufo et al. 2013, Craig et al., 2018). That said, this assumes that soils are composed of the mineral soil components suitable for stabilization of organic C (e.g. clay-rich soils) and that mineral surfaces are not already saturated with organic C (Castellano et al. 2015). Regardless, we argue that longer-term studies explicitly examining changes in specific SOM fractions are needed to assess the full magnitude of how the ‘Gadgil effect’ influences on C storage in forest soils.

### Conclusions

The growing appreciation of mycorrhizal fungi as drivers of soil biogeochemical cycles has led to the promotion of incorporating tree mycorrhizal associations as trait integrators in modeling efforts (Averill et al. 2014; Phillips et al., 2013; Sulman et al., 2017). While these efforts have been fruitful in advancing model predictions of C and nutrient cycles in terrestrial ecosystems, these classifications run the risk of obscuring the vast phylogenetic and functional diversity among EM fungi (Pellitier & Zak, 2017; Zak et al., 2019), which are potentially key in understanding effects on ecosystem processes such as the ‘Gadgil effect’ (Fernandez & Kennedy, 2016). Given the notable differences that we observed between adjacent sites that differed in both plant and fungal community composition, we caution against generalizing about the role of EM fungal suppression of OM decomposition without further examination of both mechanism and context-dependency. Instead, we advocate the implementation of high-throughput molecular approaches coupled with experiments across natural environmental gradients as well as the use of techniques that specifically track resource movement from substrates into particular microbial guilds (Nuccio et al. 2013), to improve our understanding of how EM-saprotroph interactions affect belowground forest C and N cycling.

## Acknowledgements

The authors thank J. Huggins, A. Busacker, L. Mielke, C. Daws, E. Bremer, M. Corbin, and M.L. McCormack for field and laboratory assistance. Western Ag Innovations provided a PRS Research Award to C. Fernandez and further support was provided by NSF grant (DEB-1554375) to P. Kennedy.

