## Supplemental Materials for "Decelerated carbon cycling by ectomycorrhizal fungi is controlled by substrate quality and community composition"

**Supplemental Tables**

**Table S1.** Forest site tree host properties.

|  | Oak | Pine |
| --- | --- | --- |
| DBH (cm) | 20.7 (1.76) | 11.45 (0.81) |
| Basal area (m^2^) | 3.99 (0.55) | 1.67 (0.21) |

**Table S2.** Soil pH and nutrient availability at the oak and pine forest sites. All elements are reported as mean µg 10 cm^-2^ mo^-1^ with ±SE.

|  | Oak | |  | Pine | |
| --- | --- | --- | --- | --- | --- |
|  | Mean | SE |  | Mean | SE |
| pH | 4.49 | 0.1 |  | 5.39 | 0.1 |
| NO_3_- | 45.3 | 8.7 |  | 60.7 | 15.7 |
| NH_4_+ | 6.4 | 1.1 |  | 8.0 | 2.3 |
| Ca | 591.3 | 54.9 |  | 663.7 | 54.9 |
| Mg | 170.0 | 15.4 |  | 138.6 | 9.8 |
| K | 289.4 | 51.4 |  | 150.0 | 18.1 |
| P | 13.9 | 2.1 |  | 8.0 | 0.8 |
| Fe | 3.4 | 0.5 |  | 2.0 | 0.3 |
| Mn | 19.3 | 4.1 |  | 3.1 | 0.6 |
| Zn | 1.5 | 0.2 |  | 0.6 | 0.1 |
| S | 6.1 | 0.9 |  | 3.6 | 0.5 |
| Al | 21.4 | 1.8 |  | 13.2 | 1.7 |

**Table S3.** Fungal guild assignments for fungal Genera

| **Phylum** | **Genus** | **Guild** |
| --- | --- | --- |
| Ascomycota | *Absconditella* | Other Symbiotroph |
| Ascomycota | *Acrodontium* | Pathotroph |
| Ascomycota | *Aequabiliella* | Pathotroph |
| Ascomycota | *Allantophomopsis* | Pathotroph |
| Ascomycota | *Alternaria* | Saprotroph |
| Ascomycota | *Ampelomyces* | Other Symbiotroph |
| Ascomycota | *Angustimassarina* | Saprotroph |
| Ascomycota | *Anthostomella* | Saprotroph |
| Ascomycota | *Apiospora* | Saprotroph |
| Ascomycota | *Apodus* | Saprotroph |
| Ascomycota | *Arachnopeziza* | Saprotroph |
| Ascomycota | *Archaeorhizomyces* | Saprotroph |
| Ascomycota | *Articulospora* | Saprotroph |
| Ascomycota | *Ascobolus* | Saprotroph |
| Ascomycota | *Aspergillus* | Saprotroph |
| Ascomycota | *Aureobasidium* | Pathotroph |
| Ascomycota | *Beauveria* | Pathotroph |
| Ascomycota | *Bipolaris* | Pathotroph |
| Ascomycota | *Biscogniauxia* | Saprotroph |
| Ascomycota | *Bisporella* | Saprotroph |
| Ascomycota | *Botryotinia* | Pathotroph |
| Ascomycota | *Byssonectria* | Saprotroph |
| Ascomycota | *Calcarisporium* | Pathotroph |
| Ascomycota | *Camarosporium* | Pathotroph |
| Ascomycota | *Candelaria* | Other Symbiotroph |
| Ascomycota | *Candelariella* | Other Symbiotroph |
| Ascomycota | *Capnobotryella* | Saprotroph |
| Ascomycota | *Capronia* | Other Symbiotroph |
| Ascomycota | *Cenococcum* | Ectomycorrhizal |
| Ascomycota | *Ceratocladium* | Saprotroph |
| Ascomycota | *Chaetomella* | Saprotroph |
| Ascomycota | *Chaetomium* | Saprotroph |
| Ascomycota | *Chaetosphaeria* | Saprotroph |
| Ascomycota | *Chloridium* | Ectomycorrhizal |
| Ascomycota | *Cladophialophora* | Saprotroph |
| Ascomycota | *Cladosporium* | Saprotroph |
| Ascomycota | *Clonostachys* | Pathotroph |
| Ascomycota | *Clypeosphaeria* | Saprotroph |
| Ascomycota | *Coccomyces* | Pathotroph |
| Ascomycota | *Colletotrichum* | Pathotroph |
| Ascomycota | *Collophora* | Pathotroph |
| Ascomycota | *Coniochaeta* | Saprotroph |
| Ascomycota | *Coniothyrium* | Saprotroph |
| Ascomycota | *Corynespora* | Saprotroph |
| Ascomycota | *Cryptodiscus* | Saprotroph |
| Ascomycota | *Curvularia* | Pathotroph |
| Ascomycota | *Cyberlindnera* | Mold_Yeast |
| Ascomycota | *Cylindrium* | Saprotroph |
| Ascomycota | *Cyphellophora* | Saprotroph |
| Ascomycota | *Dactylella* | Saprotroph |
| Ascomycota | *Dactylellina* | Saprotroph |
| Ascomycota | *Dactylonectria* | Saprotroph |
| Ascomycota | *Debaryomyces* | Mold_Yeast |
| Ascomycota | *Devriesia* | Pathotroph |
| Ascomycota | *Dichotomopilus* | Saprotroph |
| Ascomycota | *Dictyochaeta* | Saprotroph |
| Ascomycota | *Dictyosporium* | Saprotroph |
| Ascomycota | *Didymella* | Saprotroph |
| Ascomycota | *Dinemasporium* | Saprotroph |
| Ascomycota | *Diplodia* | Saprotroph |
| Ascomycota | *Discosia* | Pathotroph |
| Ascomycota | *Dothidea* | Saprotroph |
| Ascomycota | *Dothiorella* | Pathotroph |
| Ascomycota | *Eleutheromyces* | Unknown |
| Ascomycota | *Epicoccum* | Pathotroph |
| Ascomycota | *Eucasphaeria* | Saprotroph |
| Ascomycota | *Exophiala* | Saprotroph |
| Ascomycota | *Exserohilum* | Pathotroph |
| Ascomycota | *Fusarium* | Saprotroph |
| Ascomycota | *Fusicladium* | Pathotroph |
| Ascomycota | *Arthrocatena* | Saprotroph |
| Ascomycota | *Cadophora* | Other Symbiotroph |
| Ascomycota | *Candida* | Mold_Yeast |
| Ascomycota | *Candida* | Mold_Yeast |
| Ascomycota | *Dendrophoma* | Pathotroph |
| Ascomycota | *Mycoarthris* | Saprotroph |
| Ascomycota | *Myxozyma* | Mold_Yeast |
| Ascomycota | *Spirosphaera* | Saprotroph |
| Ascomycota | *Trichothecium* | Pathotroph |
| Ascomycota | *Xenochalara* | Saprotroph |
| Ascomycota | *Genea* | Ectomycorrhizal |
| Ascomycota | *Geomyces* | Saprotroph |
| Ascomycota | *Gorgomyces* | Unknown |
| Ascomycota | *Gymnostellatospora* | Saprotroph |
| Ascomycota | *Haptocillium* | Pathotroph |
| Ascomycota | *Helicoma* | Saprotroph |
| Ascomycota | *Helminthosporium* | Pathotroph |
| Ascomycota | *Helvella* | Ectomycorrhizal |
| Ascomycota | *Hemileucoglossum* | Unknown |
| Ascomycota | *Hendersonia* | Saprotroph |
| Ascomycota | *Herpotrichia* | Saprotroph |
| Ascomycota | *Humaria* | Ectomycorrhizal |
| Ascomycota | *Hyaloscypha* | Saprotroph |
| Ascomycota | *Hydnotrya* | Ectomycorrhizal |
| Ascomycota | *Hymenoscyphus* | Saprotroph |
| Ascomycota | *Hyphodiscosia* | Unknown |
| Ascomycota | *Hypomyces* | Saprotroph |
| Ascomycota | *Hypoxylon* | Saprotroph |
| Ascomycota | *Hysterographium* | Saprotroph |
| Ascomycota | *Ilyonectria* | Saprotroph |
| Ascomycota | *Infundichalara* | Saprotroph |
| Ascomycota | *Isaria* | Saprotroph |
| Ascomycota | *Kabatina* | Saprotroph |
| Ascomycota | *Kalmusia* | Saprotroph |
| Ascomycota | *Keissleriella* | Saprotroph |
| Ascomycota | *Knufia* | Saprotroph |
| Ascomycota | *Lachnellula* | Saprotroph |
| Ascomycota | *Lachnum* | Saprotroph |
| Ascomycota | *Latorua* | Unknown |
| Ascomycota | *Lecanicillium* | Pathotroph |
| Ascomycota | *Leotia* | Saprotroph |
| Ascomycota | *Leuconeurospora* | Saprotroph |
| Ascomycota | *Leucothecium* | Saprotroph |
| Ascomycota | *Lophiostoma* | Saprotroph |
| Ascomycota | *Lophiotrema* | Saprotroph |
| Ascomycota | *Lophium* | Saprotroph |
| Ascomycota | *Lophodermium* | Pathotroph |
| Ascomycota | *Mariannaea* | Saprotroph |
| Ascomycota | *Massarina* | Saprotroph |
| Ascomycota | *Meliniomyces* | Other Symbiotroph |
| Ascomycota | *Metacordyceps* | Pathotroph |
| Ascomycota | *Metapochonia* | Saprotroph |
| Ascomycota | *Metarhizium* | Pathotroph |
| Ascomycota | *Microcera* | Saprotroph |
| Ascomycota | *Microcyclospora* | Unknown |
| Ascomycota | *Monacrosporium* | Saprotroph |
| Ascomycota | *Monilinia* | Pathotroph |
| Ascomycota | *Monochaetia* | Pathotroph |
| Ascomycota | *Muriphaeosphaeria* | Unknown |
| Ascomycota | *Mycosymbioces* | Unknown |
| Ascomycota | *Myelochroa* | Other Symbiotroph |
| Ascomycota | *Myriangium* | Pathotroph |
| Ascomycota | *Myrmecridium* | Saprotroph |
| Ascomycota | *Myrothecium* | Saprotroph |
| Ascomycota | *Nectria* | Saprotroph |
| Ascomycota | *Nemania* | Saprotroph |
| Ascomycota | *Neocatenulostroma* | Unknown |
| Ascomycota | *Neodevriesia* | Unknown |
| Ascomycota | *Neopestalotiopsis* | Unknown |
| Ascomycota | *Nigrospora* | Saprotroph |
| Ascomycota | *Ochroconis* | Saprotroph |
| Ascomycota | *Oidiodendron* | Pathotroph |
| Ascomycota | *Ophiocordyceps* | Pathotroph |
| Ascomycota | *Ophiognomonia* | Pathotroph |
| Ascomycota | *Ophiostoma* | Pathotroph |
| Ascomycota | *Orbilia* | Saprotroph |
| Ascomycota | *Otidea* | Ectomycorrhizal |
| Ascomycota | *Pachyphlodes* | Unknown |
| Ascomycota | *Pachyphloeus* | Ectomycorrhizal |
| Ascomycota | *Paecilomyces* | Saprotroph |
| Ascomycota | *Paracamarosporium* | Saprotroph |
| Ascomycota | *Paraconiothyrium* | Saprotroph |
| Ascomycota | *Paraphaeosphaeria* | Saprotroph |
| Ascomycota | *Paraphoma* | Unknown |
| Ascomycota | *Parastagonospora* | Unknown |
| Ascomycota | *Penicillium* | Mold_Yeast |
| Ascomycota | *Periconia* | Saprotroph |
| Ascomycota | *Perusta* | Unknown |
| Ascomycota | *Pestalotiopsis* | Pathotroph |
| Ascomycota | *Pezicula* | Saprotroph |
| Ascomycota | *Peziza* | Ectomycorrhizal |
| Ascomycota | *Pezoloma* | Saprotroph |
| Ascomycota | *Phacidium* | Pathotroph |
| Ascomycota | *Phaeoacremonium* | Pathotroph |
| Ascomycota | *Phaeococcomyces* | Saprotroph |
| Ascomycota | *Phaeosphaeria* | Saprotroph |
| Ascomycota | *Phaeothecoidiella* | Unknown |
| Ascomycota | *Phialocephala* | Other Symbiotroph |
| Ascomycota | *Phialophora* | Other Symbiotroph |
| Ascomycota | *Phoma* | Pathotroph |
| Ascomycota | *Phoma* | Saprotroph |
| Ascomycota | *Phomatospora* | Saprotroph |
| Ascomycota | *Pilidiella* | Pathotroph |
| Ascomycota | *Plectania* | Saprotroph |
| Ascomycota | *Podospora* | Saprotroph |
| Ascomycota | *Preussia* | Saprotroph |
| Ascomycota | *Prosthemium* | Saprotroph |
| Ascomycota | *Pseudeurotium* | Saprotroph |
| Ascomycota | *Pseudocosmospora* | Saprotroph |
| Ascomycota | *Pseudodictyosporium* | Saprotroph |
| Ascomycota | *Pseudogymnoascus* | Saprotroph |
| Ascomycota | *Pseudopithomyces* | Saprotroph |
| Ascomycota | *Pseudoplectania* | Saprotroph |
| Ascomycota | *Pseudosigmoidea* | Unknown |
| Ascomycota | *Pustularia* | Unknown |
| Ascomycota | *Pyrenochaeta* | Saprotroph |
| Ascomycota | *Pyrenochaetopsis* | Saprotroph |
| Ascomycota | *Rachicladosporium* | Saprotroph |
| Ascomycota | *Ramularia* | Pathotroph |
| Ascomycota | *Rhinocladiella* | Saprotroph |
| Ascomycota | *Robillarda* | Unknown |
| Ascomycota | *Rosellinia* | Saprotroph |
| Ascomycota | *Rutstroemia* | Saprotroph |
| Ascomycota | *Sarocladium* | Saprotroph |
| Ascomycota | *Schizosaccharomyces* | Saprotroph |
| Ascomycota | *Scleroconidioma* | Pathotroph |
| Ascomycota | *Sclerostagonospora* | Saprotroph |
| Ascomycota | *Scolecobasidium* | Saprotroph |
| Ascomycota | *Scytalidium* | Saprotroph |
| Ascomycota | *Septorioides* | Other Symbiotroph |
| Ascomycota | *Setomelanomma* | Unknown |
| Ascomycota | *Simplicillium* | Pathotroph |
| Ascomycota | *Spathularia* | Saprotroph |
| Ascomycota | *Sphaerulina* | Other Symbiotroph |
| Ascomycota | *Sporormiella* | Saprotroph |
| Ascomycota | *Sporothrix* | Saprotroph |
| Ascomycota | *Stagonospora* | Pathotroph |
| Ascomycota | *Stagonosporopsis* | Pathotroph |
| Ascomycota | *Stemphylium* | Pathotroph |
| Ascomycota | *Stemphylium* | Saprotroph |
| Ascomycota | *Sydowia* | Saprotroph |
| Ascomycota | *Sympodiella* | Saprotroph |
| Ascomycota | *Talaromyces* | Saprotroph |
| Ascomycota | *Taphrina* | Pathotroph |
| Ascomycota | *Thelonectria* | Saprotroph |
| Ascomycota | *Tricharina* | Saprotroph |
| Ascomycota | *Trichocladium* | Saprotroph |
| Ascomycota | *Trichoderma* | Mold_Yeast |
| Ascomycota | *Trichomerium* | Other Symbiotroph |
| Ascomycota | *Tuber* | Ectomycorrhizal |
| Ascomycota | *Valsaria* | Saprotroph |
| Ascomycota | *Venturia* | Pathotroph |
| Ascomycota | *Veronaea* | Pathotroph |
| Ascomycota | *Verticillium* | Pathotroph |
| Ascomycota | *Wilcoxina* | Ectomycorrhizal |
| Ascomycota | *Yamadazyma* | Mold_Yeast |
| Ascomycota | *Zetiasplozna* | Saprotroph |
| Basidiomycota | *Agrocybe* | Saprotroph |
| Basidiomycota | *Amanita* | Ectomycorrhizal |
| Basidiomycota | *Amphinema* | Ectomycorrhizal |
| Basidiomycota | *Ampulloclitocybe* | Saprotroph |
| Basidiomycota | *Apiotrichum* | Saprotroph |
| Basidiomycota | *Armillaria* | Saprotroph |
| Basidiomycota | *Asterostroma* | Saprotroph |
| Basidiomycota | *Athelia* | Other Symbiotroph |
| Basidiomycota | *Athelia* | Saprotroph |
| Basidiomycota | *Auriculibuller* | Mold_Yeast |
| Basidiomycota | *Boletus* | Ectomycorrhizal |
| Basidiomycota | *Botryobasidium* | Saprotroph |
| Basidiomycota | *Bovista* | Saprotroph |
| Basidiomycota | *Bullera* | Mold_Yeast |
| Basidiomycota | *Bulleromyces* | Mold_Yeast |
| Basidiomycota | *Byssocorticium* | Ectomycorrhizal |
| Basidiomycota | *Carcinomyces* | Mold_Yeast |
| Basidiomycota | *Ceraceomyces* | Saprotroph |
| Basidiomycota | *Ceriporia* | Saprotroph |
| Basidiomycota | *Cerrena* | Saprotroph |
| Basidiomycota | *Clavaria* | Saprotroph |
| Basidiomycota | *Clavulina* | Ectomycorrhizal |
| Basidiomycota | *Clavulinopsis* | Saprotroph |
| Basidiomycota | *Clitocella* | Saprotroph |
| Basidiomycota | *Clitocybe* | Saprotroph |
| Basidiomycota | *Coniophora* | Saprotroph |
| Basidiomycota | *Conocybe* | Saprotroph |
| Basidiomycota | *Coprinellus* | Saprotroph |
| Basidiomycota | *Coprinus* | Saprotroph |
| Basidiomycota | *Cortinarius* | Ectomycorrhizal |
| Basidiomycota | *Cotylidia* | Unknown |
| Basidiomycota | *Craterellus* | Ectomycorrhizal |
| Basidiomycota | *Crepidotus* | Saprotroph |
| Basidiomycota | *Crinipellis* | Saprotroph |
| Basidiomycota | *Cristinia* | Saprotroph |
| Basidiomycota | *Cryptococcus* | Mold_Yeast |
| Basidiomycota | *Cryptococcus* | Mold_Yeast |
| Basidiomycota | *Cutaneotrichosporon* | Unknown |
| Basidiomycota | *Cystobasidiopsis* | Saprotroph |
| Basidiomycota | *Cystobasidium* | Pathotroph |
| Basidiomycota | *Cystodermella* | Saprotroph |
| Basidiomycota | *Cystofilobasidium* | Unknown |
| Basidiomycota | *Dioszegia* | Mold_Yeast |
| Basidiomycota | *Entoloma* | Saprotroph |
| Basidiomycota | *Erythrobasidium* | Unknown |
| Basidiomycota | *Fellomyces* | Mold_Yeast |
| Basidiomycota | *Fibulobasidium* | Mold_Yeast |
| Basidiomycota | *Filobasidium* | Saprotroph |
| Basidiomycota | *Flagelloscypha* | Saprotroph |
| Basidiomycota | *Flavodon* | Saprotroph |
| Basidiomycota | *Galzinia* | Saprotroph |
| Basidiomycota | *Geminibasidium* | Saprotroph |
| Basidiomycota | *Genolevuria* | Mold_Yeast |
| Basidiomycota | *Gloeodontia* | Saprotroph |
| Basidiomycota | *Goffeauzyma* | Unknown |
| Basidiomycota | *Guehomyces* | Unknown |
| Basidiomycota | *Gymnopus* | Saprotroph |
| Basidiomycota | *Gyroporus* | Ectomycorrhizal |
| Basidiomycota | *Hannaella* | Mold_Yeast |
| Basidiomycota | *Hebeloma* | Ectomycorrhizal |
| Basidiomycota | *Heterocephalacria* | Unknown |
| Basidiomycota | *Hortiboletus* | Saprotroph |
| Basidiomycota | *Hydnocristella* | Saprotroph |
| Basidiomycota | *Hydropus* | Saprotroph |
| Basidiomycota | *Hygrophorus* | Ectomycorrhizal |
| Basidiomycota | *Hyphoderma* | Saprotroph |
| Basidiomycota | *Hyphodermella* | Saprotroph |
| Basidiomycota | *Hyphodontia* | Saprotroph |
| Basidiomycota | *Inocybe* | Ectomycorrhizal |
| Basidiomycota | *Irpex* | Saprotroph |
| Basidiomycota | *Ischnoderma* | Saprotroph |
| Basidiomycota | *Kavinia* | Saprotroph |
| Basidiomycota | *Kockovaella* | Mold_Yeast |
| Basidiomycota | *Krasilnikovozyma* | Unknown |
| Basidiomycota | *Kwoniella* | Mold_Yeast |
| Basidiomycota | *Laccaria* | Ectomycorrhizal |
| Basidiomycota | *Lactarius* | Ectomycorrhizal |
| Basidiomycota | *Lentinus* | Saprotroph |
| Basidiomycota | *Lepiota* | Saprotroph |
| Basidiomycota | *Lepista* | Saprotroph |
| Basidiomycota | *Leucocybe* | Unknown |
| Basidiomycota | *Leucosporidium* | Unknown |
| Basidiomycota | *Limacella* | Saprotroph |
| Basidiomycota | *Luellia* | Saprotroph |
| Basidiomycota | *Lycoperdon* | Saprotroph |
| Basidiomycota | *Marasmiellus* | Saprotroph |
| Basidiomycota | *Marasmius* | Saprotroph |
| Basidiomycota | *Membranomyces* | Ectomycorrhizal |
| Basidiomycota | *Minimedusa* | Unknown |
| Basidiomycota | *Mutinus* | Saprotroph |
| Basidiomycota | *Mycena* | Saprotroph |
| Basidiomycota | *Mycenella* | Unknown |
| Basidiomycota | *Naganishia* | Unknown |
| Basidiomycota | *Oberwinklerozyma* | Unknown |
| Basidiomycota | *Oberwinklerozyma* | Unknown |
| Basidiomycota | *Oberwinklerozyma* | Unknown |
| Basidiomycota | *Occultifur* | Unknown |
| Basidiomycota | *Odontia* | Ectomycorrhizal |
| Basidiomycota | *Paralepista* | Unknown |
| Basidiomycota | *Paxillus* | Ectomycorrhizal |
| Basidiomycota | *Peniophora* | Saprotroph |
| Basidiomycota | *Perenniporia* | Saprotroph |
| Basidiomycota | *Phaeoclavulina* | Ectomycorrhizal |
| Basidiomycota | *Phaeotremella* | Mold_Yeast |
| Basidiomycota | *Phanerochaete* | Saprotroph |
| Basidiomycota | *Phenoliferia* | Unknown |
| Basidiomycota | *Pholiota* | Saprotroph |
| Basidiomycota | *Piloderma* | Ectomycorrhizal |
| Basidiomycota | *Piskurozyma* | Unknown |
| Basidiomycota | *Pluteus* | Saprotroph |
| Basidiomycota | *Postia* | Saprotroph |
| Basidiomycota | *Psathyrella* | Saprotroph |
| Basidiomycota | *Pseudoclitocybe* | Saprotroph |
| Basidiomycota | *Pseudohyphozyma* | Unknown |
| Basidiomycota | *Pseudohyphozyma* | Unknown |
| Basidiomycota | *Pseudotomentella* | Ectomycorrhizal |
| Basidiomycota | *Ramaria* | Ectomycorrhizal |
| Basidiomycota | *Rectipilus* | Saprotroph |
| Basidiomycota | *Rhizoctonia* | Saprotroph |
| Basidiomycota | *Rhizocybe* | Unknown |
| Basidiomycota | *Rhodocollybia* | Saprotroph |
| Basidiomycota | *Rhodosporidiobolus* | Unknown |
| Basidiomycota | *Rhodotorula* | Saprotroph |
| Basidiomycota | *Ripartites* | Saprotroph |
| Basidiomycota | *Russula* | Ectomycorrhizal |
| Basidiomycota | *Saitozyma* | Mold_Yeast |
| Basidiomycota | *Sarcoporia* | Saprotroph |
| Basidiomycota | *Scleroderma* | Ectomycorrhizal |
| Basidiomycota | *Scytinostromella* | Saprotroph |
| Basidiomycota | *Sebacina* | Other Symbiotroph |
| Basidiomycota | *Serendipita* | Other Symbiotroph |
| Basidiomycota | *Serpula* | Saprotroph |
| Basidiomycota | *Singerocybe* | Unknown |
| Basidiomycota | *Slooffia* | Unknown |
| Basidiomycota | *Slooffia* | Unknown |
| Basidiomycota | *Solicoccozyma* | Unknown |
| Basidiomycota | *Sporobolomyces* | Saprotroph |
| Basidiomycota | *Steccherinum* | Saprotroph |
| Basidiomycota | *Stereum* | Saprotroph |
| Basidiomycota | *Strobilomyces* | Ectomycorrhizal |
| Basidiomycota | *Suillus* | Ectomycorrhizal |
| Basidiomycota | *Suillus* | Ectomycorrhizal |
| Basidiomycota | *Syzygospora* | Mold_Yeast |
| Basidiomycota | *Thelephora* | Ectomycorrhizal |
| Basidiomycota | *Tomentella* | Ectomycorrhizal |
| Basidiomycota | *Trametes* | Saprotroph |
| Basidiomycota | *Trechispora* | Saprotroph |
| Basidiomycota | *Tremella* | Mold_Yeast |
| Basidiomycota | *Tricholoma* | Ectomycorrhizal |
| Basidiomycota | *Tritirachium* | Saprotroph |
| Basidiomycota | *Tylopilus* | Ectomycorrhizal |
| Basidiomycota | *Tylospora* | Ectomycorrhizal |
| Basidiomycota | *Udeniomyces* | Unknown |
| Basidiomycota | *Vanrija* | Unknown |
| Basidiomycota | *Vararia* | Saprotroph |
| Basidiomycota | *Vishniacozyma* | Mold_Yeast |
| Basidiomycota | *Volvariella* | Saprotroph |
| Basidiomycota | *Wallemia* | Saprotroph |
| Basidiomycota | *Xenasmatella* | Saprotroph |
| Basidiomycota | *Xerocomus* | Ectomycorrhizal |
| Basidiomycota | *Yamadamyces* | Unknown |
| Basidiomycota | *Yurkovia* | Unknown |
| Calcarisporiellomycota | *Calcarisporiella* | Unknown |
| Entomophthoromycota | *Basidiobolus* | Saprotroph |
| Glomeromycota | *Archaeospora* | Other Symbiotroph |
| Glomeromycota | *Glomus* | Other Symbiotroph |
| Glomeromycota | *Paraglomus* | Other Symbiotroph |
| Glomeromycota | *Rhizophagus* | Other Symbiotroph |
| Kickxellomycota | *Spiromyces* | Saprotroph |
| Mortierellomycota | *Mortierella* | Mold_Yeast |
| Mucoromycota | *Backusella* | Mold_Yeast |
| Mucoromycota | *Mucor* | Mold_Yeast |
| Mucoromycota | *Rhizopus* | Mold_Yeast |
| Mucoromycota | *Umbelopsis* | Mold_Yeast |

**Table S4.** Mixed model effects tests for litter chemistry after 12 month incubation.

|  |  |  |  | Nitrogen | |  |  | Carbon | |  |  | Lignin | |  |  | Cellulose | |  |  | Hemicellulose | |  |
| --- | --- | --- | --- | --- | --- | --- | --- | --- | --- | --- | --- | --- | --- | --- | --- | --- | --- | --- | --- | --- | --- | --- |
| Fixed Effects | dfnum | dfden |  | *F* | *P* |  |  | *F* | *P* |  |  | *F* | *P* |  |  | *F* | *P* |  |  | *F* | *P* |  |
| (Intercept) | 1 | 28 |  | 2056.10 | <.0001 | *** |  | 8509.39 | <.0001 | *** |  | 1138.01 | <.0001 | *** |  | 577.45 | <.0001 | *** |  | 564.01 | <.0001 | *** |
| Litter Type (L) | 1 | 28 |  | 83.85 | <.0001 | *** |  | 8.28 | 0.01 | ** |  | 1.40 | 0.25 |  |  | 9.11 | 0.01 | * |  | 1.61 | 0.22 |  |
| Stand (S) | 1 | 10 |  | 2.12 | 0.18 |  |  | 5.99 | 0.03 | * |  | 6.76 | 0.03 | * |  | 0.57 | 0.47 |  |  | 0.01 | 0.94 |  |
| Trenching (T) | 1 | 28 |  | 0.86 | 0.36 |  |  | 5.23 | 0.03 | * |  | 2.80 | 0.11 |  |  | 7.84 | 0.01 | * |  | 8.06 | 0.01 | * |
| L × S | 1 | 28 |  | 7.63 | 0.01 | * |  | 0.01 | 0.92 |  |  | 0.02 | 0.89 |  |  | 0.17 | 0.69 |  |  | 0.38 | 0.54 |  |
| L × T | 1 | 28 |  | 0.05 | 0.82 |  |  | 0.35 | 0.56 |  |  | 5.71 | 0.02 | * |  | 0.19 | 0.66 |  |  | 0.04 | 0.84 |  |
| S × T | 1 | 28 |  | 0.08 | 0.78 |  |  | 7.30 | 0.01 | ** |  | 3.33 | 0.08 |  |  | 0.18 | 0.68 |  |  | 0.76 | 0.39 |  |
| L × S × T | 1 | 28 |  | 0.02 | 0.88 |  |  | 1.84 | 0.19 |  |  | 5.14 | 0.03 | * |  | 0.60 | 0.44 |  |  | 2.68 | 0.11 |  |

**Table S5.** ANCOVA model

|  |  |  |  |  | Carbon content | |  |  | Lignin content | |  |
| --- | --- | --- | --- | --- | --- | --- | --- | --- | --- | --- | --- |
| *A. Oak litter* | Fixed Effects | numDF | denDF |  | *F* | *P* |  |  | *F* | *P* |  |
|  | (Intercept) | 1 | 10 |  | 1089.18 | <.0001 | *** |  | 420.5947 | <.0001 | *** |
|  | Nitrogen content (N) | 1 | 4 |  | 101.2084 | 0.0005 | ** |  | 30.8058 | 0.0052 | ** |
|  | Stand (S) | 1 | 10 |  | 4.1704 | 0.0684 |  |  | 6.4854 | 0.029 |  |
|  | Trenching (T) | 1 | 4 |  | 1.9963 | 0.2306 |  |  | 1.488 | 0.2895 |  |
|  | N x S | 1 | 4 |  | 5.3554 | 0.0817 |  |  | 0.4291 | 0.5482 |  |
|  | N x T | 1 | 4 |  | 0.111 | 0.7558 |  |  | 7.4154 | 0.0528 |  |
|  | S x T | 1 | 4 |  | 0.8756 | 0.4024 |  |  | 0.5871 | 0.4862 |  |
|  | N x S x T | 1 | 4 |  | 0.2494 | 0.6437 |  |  | 0.6151 | 0.4767 |  |
|  |  |  |  |  | Carbon content | |  |  | Lignin content | |  |
| *B. Pine litter* | Fixed Effects | numDF | denDF |  | *F* | *P* |  |  | *F* | *P* |  |
|  | (Intercept) | 1 | 10 |  | 1385.1915 | <.0001 | *** |  | 402.5779 | <.0001 | *** |
|  | Nitrogen content (N) | 1 | 6 |  | 23.9567 | 0.0027 | ** |  | 7.004 | 0.0382 | * |
|  | Stand (S) | 1 | 10 |  | 2.8411 | 0.1228 |  |  | 1.0041 | 0.34 |  |
|  | Trenching (T) | 1 | 6 |  | 10.0363 | 0.0194 | * |  | 9.5914 | 0.0212 | * |
|  | N x S | 1 | 6 |  | 0.7343 | 0.4244 |  |  | 0.5833 | 0.474 |  |
|  | N x T | 1 | 6 |  | 8.4487 | 0.0271 | * |  | 6.9362 | 0.0389 | * |
|  | S x T | 1 | 6 |  | 0.66 | 0.4476 |  |  | 0.45 | 0.5273 |  |
|  | N x S x T | 1 | 6 |  | 1.8949 | 0.2178 |  |  | 0.1544 | 0.7079 |  |

**Table S6.** Mixed model results testing the effects of on litter guild abundance after 12 months of incubation

|  |  |  |  | Ectomycorrhizal | |  |  | Saprotrophic | |  |  | Molds & Yeasts | |  |  | Pathotrophic | |  |  | Other Symbiotrophic | |  |
| --- | --- | --- | --- | --- | --- | --- | --- | --- | --- | --- | --- | --- | --- | --- | --- | --- | --- | --- | --- | --- | --- | --- |
| Fixed Effects | numDF | denDF |  | *F* | *P* |  |  | *F* | *P* |  |  | *F* | *P* |  |  | *F* | *P* |  |  | *F* | *P* |  |
| (Intercept) | 1 | 159 |  | 114.886 | <0.0001 | *** |  | 347.248 | <0.0001 | *** |  | 347.248 | <0.0001 | *** |  | 238.529 | <0.0001 | *** |  | 131.543 | <0.0001 | *** |
| Incubation (I) | 1 | 159 |  | 39.159 | <0.0001 | *** |  | 13.370 | 0.000 | ** |  | 13.370 | 0.000 | ** |  | 1.015 | 0.315 |  |  | 41.922 | <0.0001 | *** |
| Litter type (L) | 1 | 159 |  | 12.331 | 0.001 | ** |  | 2.256 | 0.135 |  |  | 2.256 | 0.135 |  |  | 33.466 | <0.0001 | *** |  | 3.301 | 0.071 |  |
| Stand (S) | 1 | 10 |  | 0.233 | 0.639 |  |  | 3.249 | 0.102 |  |  | 3.249 | 0.102 |  |  | 18.727 | 0.002 | ** |  | 0.469 | 0.509 |  |
| Trenching (T) | 1 | 159 |  | 2.333 | 0.129 |  |  | 0.001 | 0.973 |  |  | 0.001 | 0.973 |  |  | 0.119 | 0.730 |  |  | 5.450 | 0.021 | * |
| I x L | 1 | 159 |  | 3.419 | 0.066 |  |  | 0.054 | 0.817 |  |  | 0.054 | 0.817 |  |  | 1.160 | 0.283 |  |  | 13.776 | 0.000 | ** |
| I x S | 1 | 159 |  | 0.610 | 0.436 |  |  | 3.195 | 0.076 |  |  | 3.195 | 0.076 |  |  | 0.307 | 0.581 |  |  | 7.571 | 0.007 | ** |
| L x S | 1 | 159 |  | 1.536 | 0.217 |  |  | 0.002 | 0.969 |  |  | 0.002 | 0.969 |  |  | 8.313 | 0.005 | ** |  | 5.564 | 0.020 |  |
| I x T | 1 | 159 |  | 1.491 | 0.224 |  |  | 0.000 | 0.983 |  |  | 0.000 | 0.983 |  |  | 0.016 | 0.900 |  |  | 3.210 | 0.075 |  |
| L x T | 1 | 159 |  | 0.906 | 0.343 |  |  | 0.197 | 0.658 |  |  | 0.197 | 0.658 |  |  | 0.811 | 0.369 |  |  | 0.008 | 0.929 |  |
| S x T | 1 | 159 |  | 1.201 | 0.275 |  |  | 0.093 | 0.761 |  |  | 0.093 | 0.761 |  |  | 0.111 | 0.740 |  |  | 7.491 | 0.007 | ** |
| I x L x S | 1 | 159 |  | 2.654 | 0.105 |  |  | 0.045 | 0.832 |  |  | 0.045 | 0.832 |  |  | 0.607 | 0.437 |  |  | 2.441 | 0.120 |  |
| I x L x T | 1 | 159 |  | 4.123 | 0.044 | * |  | 1.292 | 0.257 |  |  | 1.292 | 0.257 |  |  | 0.148 | 0.701 |  |  | 0.946 | 0.332 |  |
| I x S x T | 1 | 159 |  | 1.091 | 0.298 |  |  | 0.219 | 0.640 |  |  | 0.219 | 0.640 |  |  | 0.164 | 0.686 |  |  | 2.960 | 0.087 |  |
| L x S x T | 1 | 159 |  | 0.452 | 0.502 |  |  | 0.002 | 0.965 |  |  | 0.002 | 0.965 |  |  | 0.994 | 0.320 |  |  | 0.300 | 0.585 |  |
| I x L x S x T | 1 | 159 |  | 0.003 | 0.958 |  |  | 0.002 | 0.968 |  |  | 0.002 | 0.968 |  |  | 0.003 | 0.955 |  |  | 8.998 | 0.003 | ** |

**Supplemental Figures**

**Figure S1.** The relationship between ectomycorrhizal root in-growth and soil fungal guild abundances at the pine and oak sites. Guild abundances are Hellinger transformed abundance.

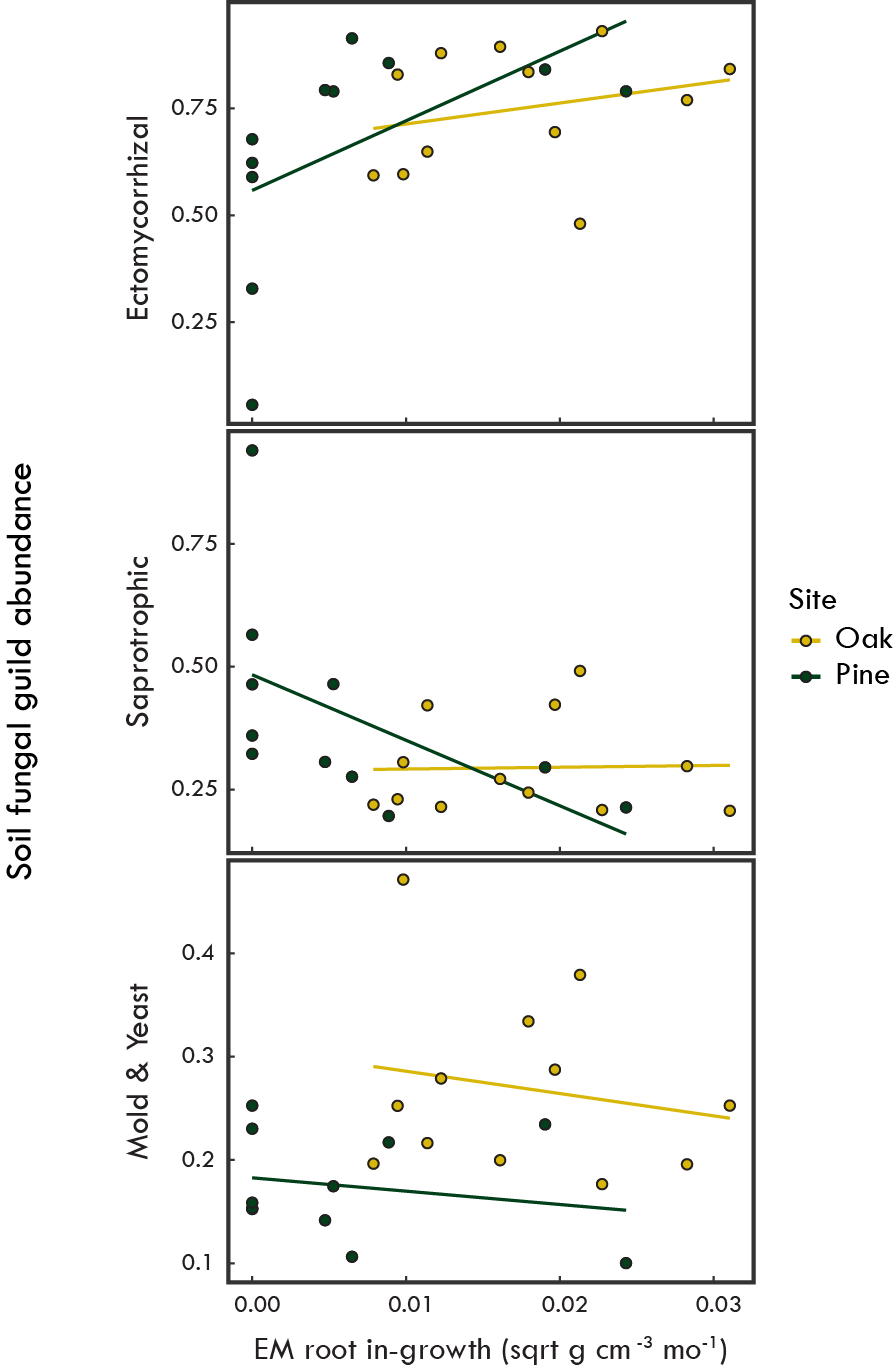

**Figure S2**. Response of root in-growth rate to the untrenched (orange) and trenched (gray) treatments at the oak and pine site.

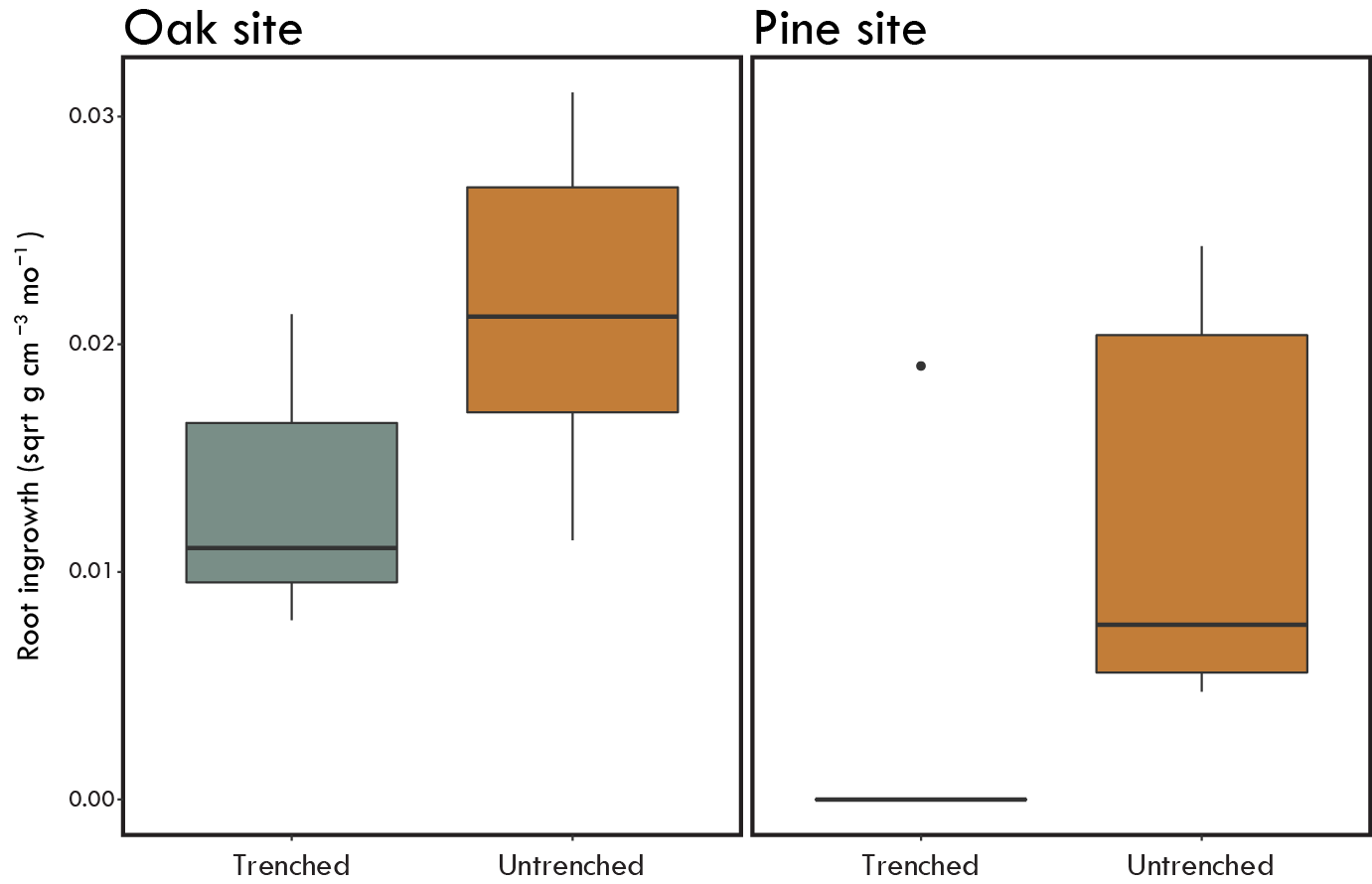

**Figure S3.** Litter mass loss results from Experiment 1. Mean percent mass remaining (±SE) of oak (A) and pine (C) litter incubated in the untrenched (orange) and trenched (gray) treatments for 2, 4 and 12 months in the corresponding forest type.

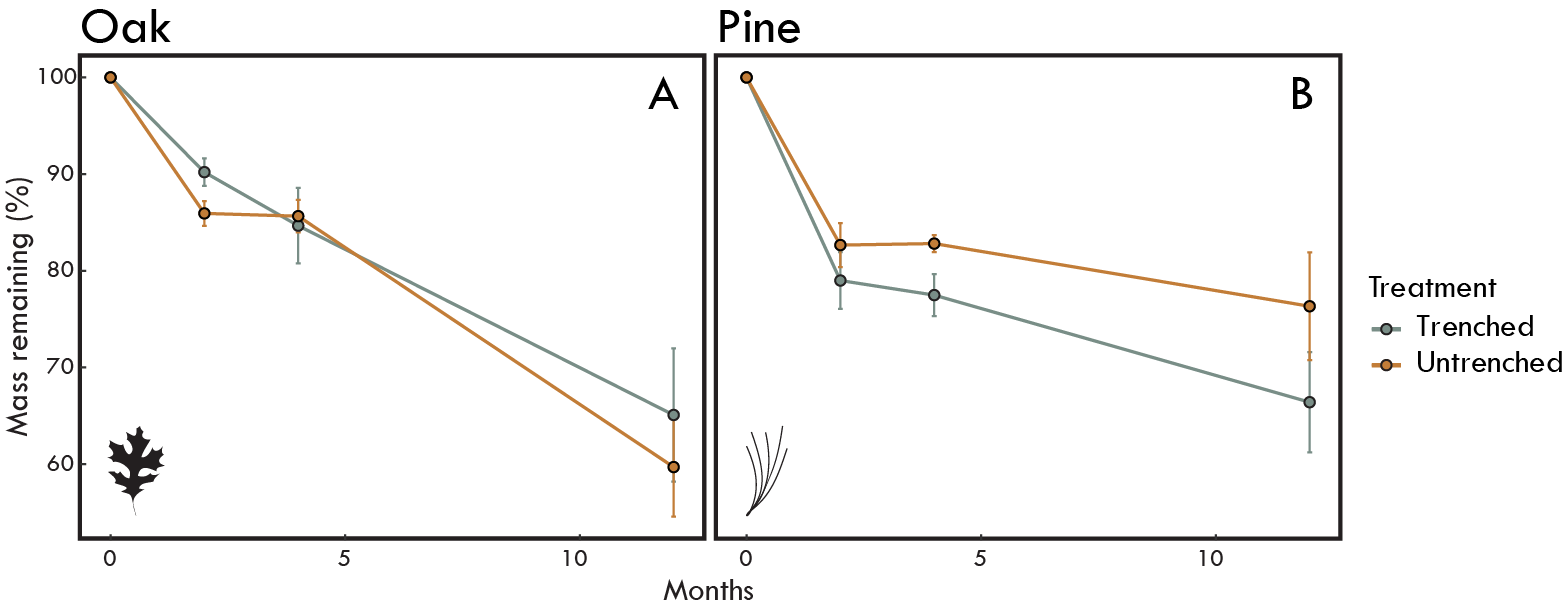

**Figure S4.** Mean abundance (Hellinger transformed) (±SE) of fungal guilds over time colonizing oak and pine litters incubated in each trenching treatment in the oak site (A) and pine site (B) in Experiment 2. Data points and lines are colored by guild.

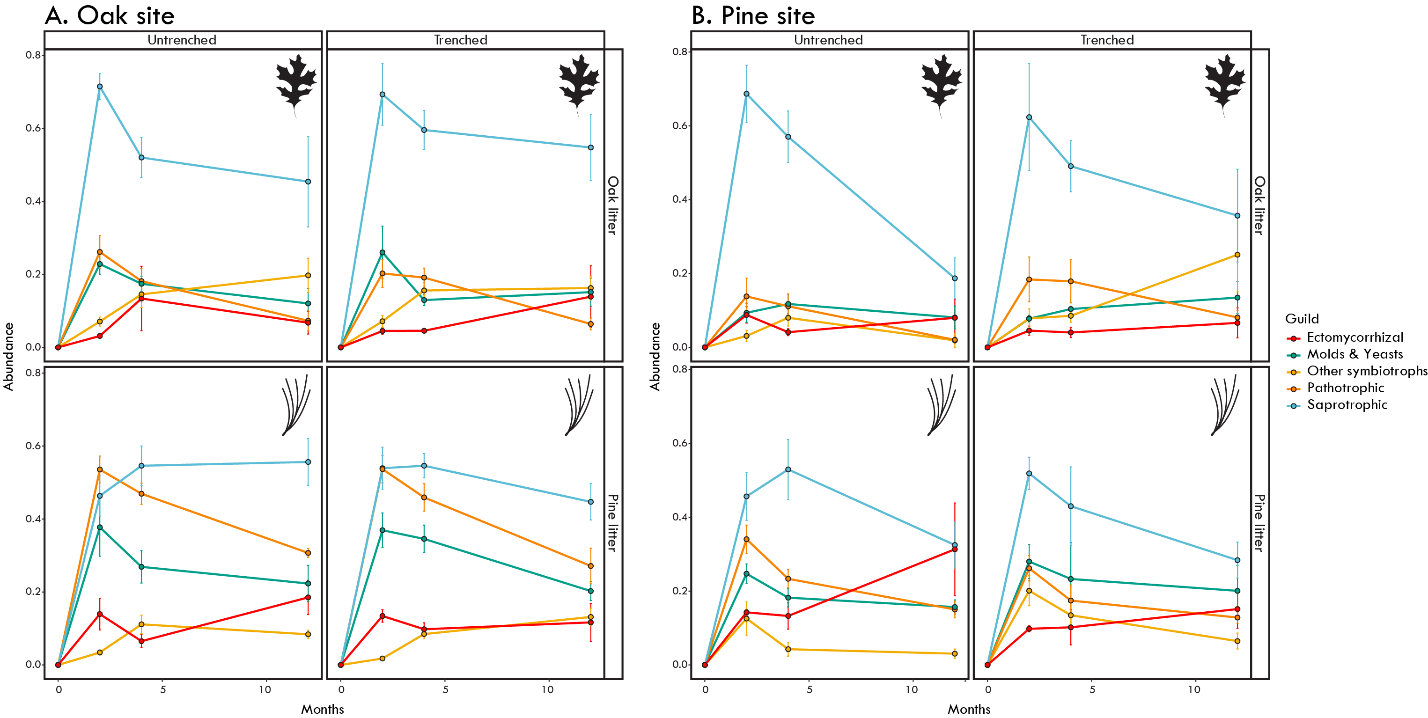

**Figure S5.** The 25 most abundant fungal genera colonizing oak and pine litters incubated in each trenching treatment in oak and pine sites. Boxplots are colored by trenching treatment.

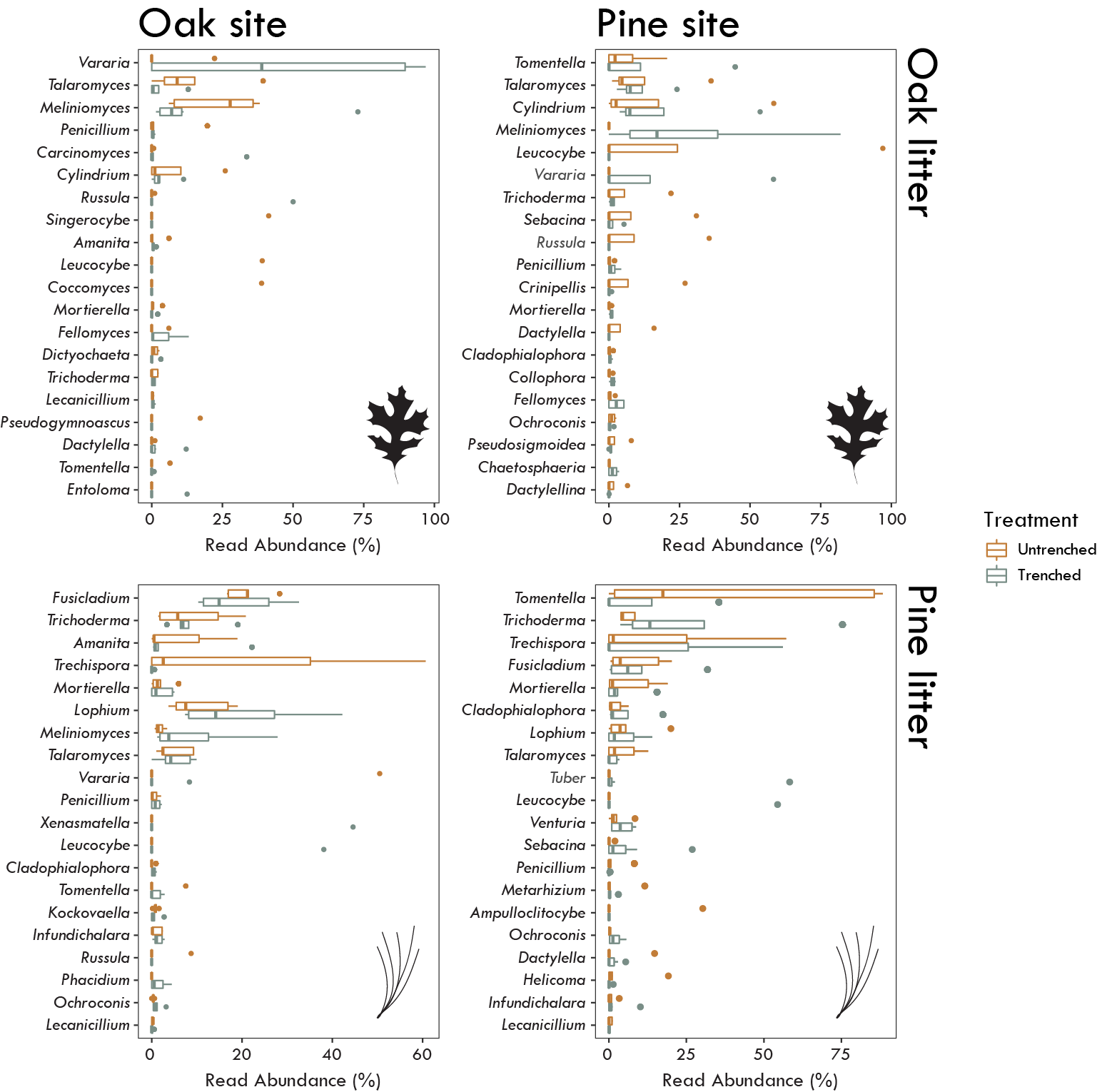

**Figure S6.** Nitrogen, carbon, and lignin concentrations (%) oak and pine litter incubated in the untrenched (orange) and trenched (gray) treatments in oak and pine sites.

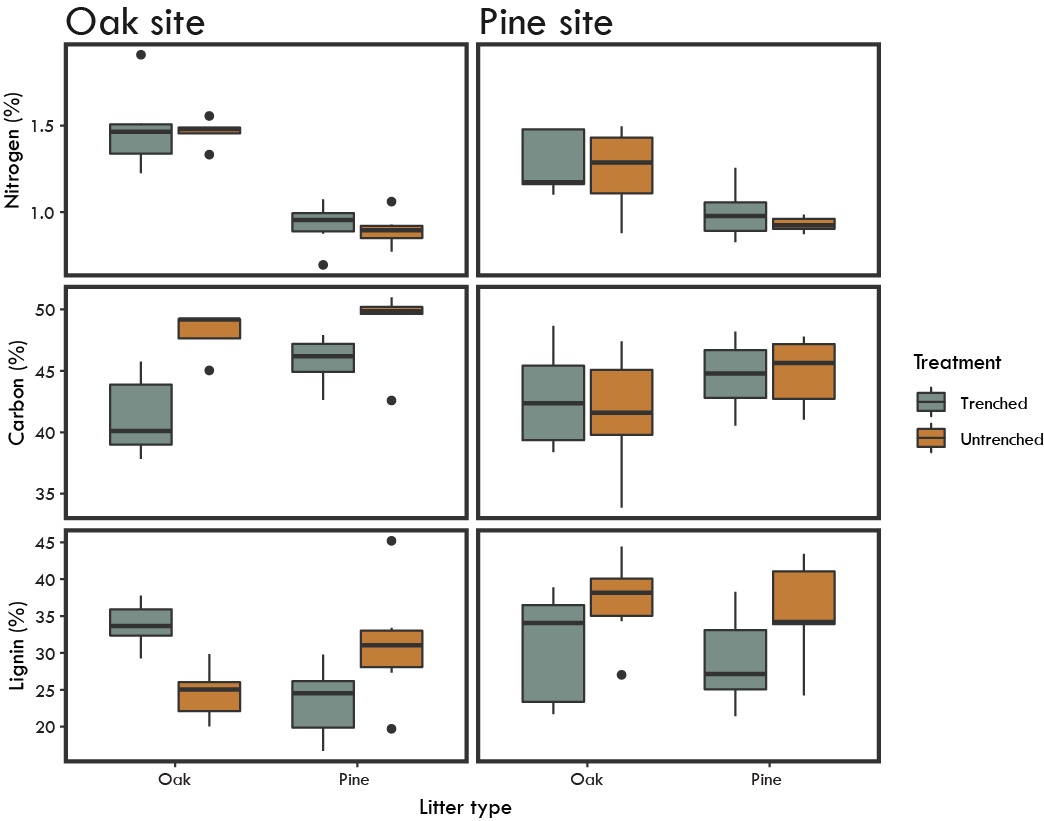
